# SAGA histone acetyltransferase module facilitates chromatin accessibility to SMC5/6

**DOI:** 10.1101/2022.09.24.508842

**Authors:** L. Mahrik, B. Stefanovie, A. Maresova, J. Princova, P. Kolesar, E. Lelkes, D. Helmlinger, M. Prevorovsky, J.J. Palecek

## Abstract

Structural Maintenance of Chromosomes (SMC) complexes are molecular machines driving chromatin organization at higher levels. In eukaryotes, three SMC complexes (cohesin, condensin, and SMC5/6) play key roles in cohesion, condensation, replication, transcription and DNA repair. Here, we performed a genetic screen in fission yeast to identify novel factors required for SMC5/6 binding to DNA. We identified 79 genes of which histone acetyltransferases (HATs) were the most represented. Genetic and phenotypic analyses suggested a particularly strong functional relationship between the SMC5/6 and SAGA complexes. Furthermore, several SMC5/6 subunits physically interacted with SAGA HAT module components Gcn5 and Ada2. As Gcn5-dependent acetylation facilitates the accessibility of chromatin to DNA repair proteins, we first analysed the formation of DNA damage-induced SMC5/6 foci in the Δ*gcn5* mutant. The SMC5/6 foci formed normally in Δ*gcn5*, suggesting SAGA-independent SMC5/6 localization to DNA-damaged sites. Next, we used Nse4-FLAG chromatin-immunoprecipitation (ChIP-seq) analysis in unchallenged cells to assess SMC5/6 distribution. A significant portion of SMC5/6 accumulated within gene regions in wild-type cells, which was reduced in Δ*gcn5* and Δ*ada2* mutants. The drop in SMC5/6 levels was also observed in *gcn5*-E191Q acetyltransferase-dead mutant. Altogether, our data suggest that the SAGA HAT module may facilitate chromatin accessibility to SMC5/6 at gene regions.

**Author Summary:** Genomes of all eukaryotes must be folded and packed into their relatively small nuclear spaces. Histones first pack free genomic DNA into nucleosomes and their arrays. Other complexes like histone modifiers and remodelers can regulate nucleosome positions and their packing within chromatin fibres. They assist in the relative opening or condensation of chromatin fibres and facilitate their accessibility to DNA-binding proteins. The highly conserved Structural Maintenance of Chromosomes (SMC) complexes (cohesin, condensin, and SMC5/6) compact further chromatin fibres at higher levels. These molecular machines can loop chromatin fibres, which need access to segments of free DNA for their physical binding to DNA. Here, we studied genetic and physical interactions between histone-modifying SAGA complex and SMC5/6. We show that the SAGA histone acetyltransferase module may facilitate chromatin access to SMC5/6.

## Introduction

Chromatin is composed of DNA and protein complexes structured at multiple levels to ensure its spatial and functional organization [1]. Histone proteins pack DNA into nucleosomes and their arrays at the basic level. At the higher levels, Structural Maintenance of Chromosomes (SMC) complexes (cohesin, condensin and SMC5/6) assist in the formation of higher-order structures like topologically associated domains or condensed mitotic chromosomes [2]. Chromatin compaction affects DNA accessibility at each level. Histone chaperones, modifiers, and remodelers can loosen, move or remodel nucleosomes to modulate essential processes like transcription or DNA repair [3, 4]. For example, the histone-modifying SAGA complex acetylates H3 histones at promoter regions, contributing to chromatin opening and facilitating the assembly of transcription initiation complexes onto core promoters and the recruitment of factors that directly interact with DNA [5].

The SMC complexes play roles in all key chromatin processes, including cohesion, condensation, replication, transcription, and DNA repair. Their cores comprise the long-armed SMC, kleisin, and kleisin-associated (KITE or HAWK) subunits [6, 7]. Uniquely, SMC5/6 complexes contain the highly conserved Nse1 and Nse2 ubiquitin- and SUMO-ligases, respectively [8, 9]. The Smc subunits are primarily built of head ATPase domains, long anti-parallel coiled-coil arms, and hinges [10–12]. Two Smc molecules form stable dimers via their hinge domains, and without ATP, their arms align into rod-like structures [13, 14]. The binding of ATP molecules to the ATPase head domains promotes the formation of large annular structures [15, 16]. The ATP binding-hydrolysis cycle drives ring-to-rod dynamic changes and promotes DNA translocation or loop extrusion [17–20].

SMC complexes were believed to interact only topologically with chromatin fibres via their large annular structures, which can embrace and traverse large chromatin complexes, including nucleosomes [12]. However, growing evidence suggests their requirement for open chromatin and direct physical binding to DNA [21–23]. Moreover, Piazza et al. [22] described the preferential binding of condensin kleisin-associated HAWK subunits to free DNA over nucleosomal DNA. Recent cryoEM analyses showed the formation of K-compartments within all SMC complexes, consisting of ATP-bound Smc heads, kleisin, and kleisin-associated subunits, which can accommodate only free DNA [16, 24–26]. In line with these findings, the SAGA complex assists in loading condensin at open chromatin regions of highly transcribed genes in fission yeast [23]. Similarly, the RSC chromatin remodelling complex recruits the Scc2-Scc4 factor to nucleosome-free regions, assisting in cohesin loading at these sites [27–29].

Recently, it was shown that the SMC5/6 K-compartment, composed of Smc5-Smc6 heads, Nse4 kleisin, and Nse1-Nse3 kleisin-associated KITE subunits, binds free DNA [16, 21]. In our previous study, we described the essential role of Nse3-DNA interaction for SMC5/6 loading or accumulation [21]. Here, we performed a genetic screen with a *nse3-R254E* fission yeast mutant that exhibits reduced DNA-binding affinity to identify new factors required for its viability. We found strong genetic interactions with the SAGA and NuA4 histone acetyltransferase (HAT) complexes. Using chromatin-immunoprecipitation (ChIP-seq) analysis, we observed a significant portion of SMC5/6 accumulated within gene regions, which was reduced in SAGA HAT deletion (Δ*ada2* and Δ*gcn5*) mutants. The magnitude of the decrease in SMC5/6 occupancy correlated with the SAGA-modified H3K9ac levels around the transcription start sites. The SMC5/6 reduced levels were also observed in *gcn5*-*E191Q* acetyltransferase-dead mutant, suggesting that the SAGA HAT module may facilitate the accessibility of chromatin for SMC5/6 loading.

## Results

### Genetic screen with a DNA-binding defective allele of the SMC5/6 complex

To identify factors that facilitate the loading or accumulation of the SMC5/6 complex on chromatin, we performed a genetic search for genes affecting the survival of cells with compromised DNA-binding ability of the SMC5/6 complex [21]. First, we created a query fission yeast strain with the DNA-binding defective *nse3-R254E* mutation in the PEM2 background (Suppl. Fig. S1A; [30]) and crossed it against the whole gene deletion yeast collection from BIONEER [31]. Using yeast colony size phenotypic readout [32], we identified 79 deletion strains that exhibited a negative genetic interaction with *nse3-R254E* (Suppl. Table ST1).

To validate our results, we randomly selected 19 of the 79 strains and crossed them with the original *nse3-R254E* strain [21]. Tetrad analysis confirmed that these mutations are synthetically sick or lethal with the *nse3-R254E* mutation (Suppl Fig. S1B). Analysis of all 79 interacting genes for the Biological process category using Gene ontologies (GO; [33]) showed the highest scores for DNA repair, chromatin organization, meiosis, and replication processes (Fig. 1A and Suppl Table ST2), in line with previous studies (reviewed in [34, 35]). Reassuringly, several *nse3-R254E* genetic interactions overlapped with the genetic interactions of other *smc5/6* mutants [21, 36–41], supporting the validity of our screen results.

**Figure 1.**
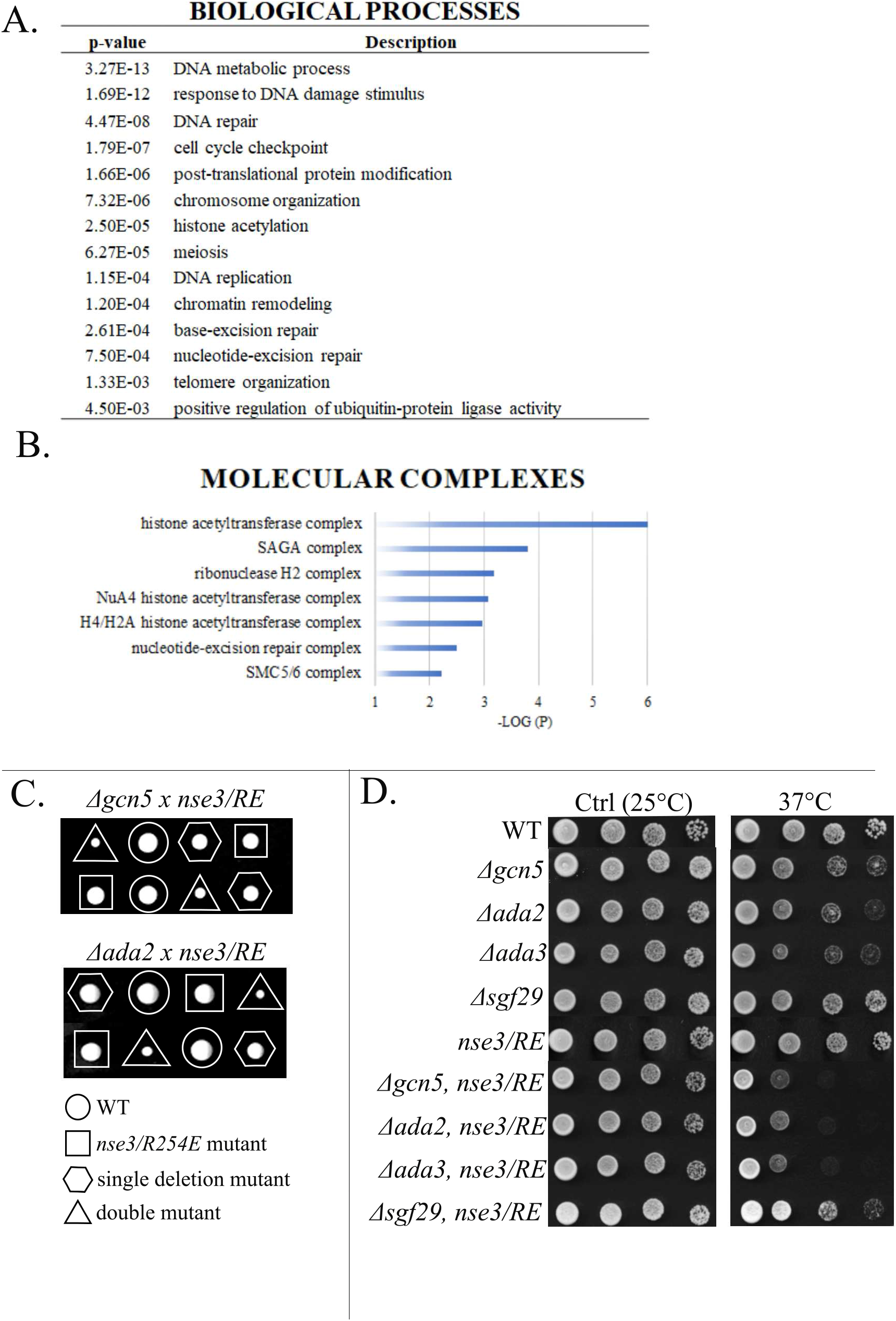
Strong genetic relation between the SMC5/6 and histone acetyltransferases. (**A**) Summary of significantly enriched GO categories for Biological processes of genes with genetic interactions with *nse3*-*R254E* (P < 0.005). (**B**) Negative log10 (P-values) evaluating the significance of the main GO Cellular component terms identified in the set of the 79 genes. Only the molecular complexes are shown. (**C**) Tetrad analysis of the heterozygous diploid fission yeast strains. The colony size of the *nse3-R254E*, Δ*gcn5* and *nse3-R254E*, Δ*ada2* double mutant is significantly reduced (triangle). Single and double mutant alleles are indicated. (**D**) Ten-fold serial dilutions of the indicated strains were plated onto YES media and grown at 25°C (control) or 37°C. The *nse3-R254E* (*nse3/RE*) mutation enhanced the sensitivity of the Δ*ada2*, Δ*ada3*, and Δ*gcn5* mutants to the higher temperature.

### Genetic interactions between SMC5/6 and histone acetyltransferase complexes

The analysis of the Cellular components GO of the 79 interacting genes showed the highest scores for histone acetyltransferase (HAT) complexes (SAGA and NuA4; Fig. 1B and Suppl Table ST3). Three (Ada2, Ada3/Ngg1, Gcn5) out of four HAT module subunits and one (Ubp8) out of four DUB module subunits of the SAGA complex were identified as hits in our screen [5]. We verified their genetic interactions using tetrad analysis (Figs. 1C and S1B) and also tested the other non-essential SAGA subunits not detected in our screen. The mating defects of the Δ*ada1*, Δ*spt7*, Δ*spt8*, and Δ*spt20* deletion mutants hampered the double mutant preparation [42]. However, the tetrad analysis of the other non-essential SAGA subunits (Suppl. Fig. S1C) showed negative genetic interactions with the *nse3-R254E* mutation (Suppl. Fig. S1D), suggesting a strong functional relationship between the SMC5/6 and SAGA complexes.

Interestingly, the temperature-sensitive (*ts*) phenotypes of the SAGA HAT module Δ*ada2*, Δ*ada3*, and Δ*gcn5* mutants were enhanced by *nse3-R254E* (Fig. 1D; [42]). The *ts* phenotypes were also enhanced by the *smc6-74* and *smc6-X* hypomorphic mutations (Suppl. Fig. S2A; [37, 43]). We also used *nse1-R188E* and *nse2-SA* mutations, which specifically abrogate DNA-repair function (ubiquitin- and SUMO-ligase activity, respectively), but not the SMC5/6 essential function [8, 44]. However, these mutations did not affect the *ts* phenotype of Δ*gcn5* (Suppl. Fig. S2A; not shown). These data suggest that the SMC5/6 essential function supports cell survival at higher temperatures in the absence of the HAT module and, conversely, that the viability of the *nse3-R254E* mutant is compromised in the absence of a functional SAGA HAT module.

### SMC5/6 and SAGA physically interact

The strong genetic relationship between the SAGA HAT module and the SMC5/6 complex prompted us to test their mutual physical interactions. First, we performed co-immunoprecipitation of Gcn5-myc and Nse4-FLAG kleisin subunit to show the association between the SAGA and SMC5/6 complexes *in vivo* (Fig. 2A). The fission yeast cells carrying Nse4-FLAG (with or without Gcn5-myc) were lysed, and proteins were precipitated with an anti-myc antibody. A small amount of Nse4-FLAG was specifically recovered in the Gcn5-myc precipitates but not in the control experiment without myc-tagged Gcn5, suggesting a weak or transient association between SAGA and SMC5/6 in the yeast cells.

**Figure 2.**
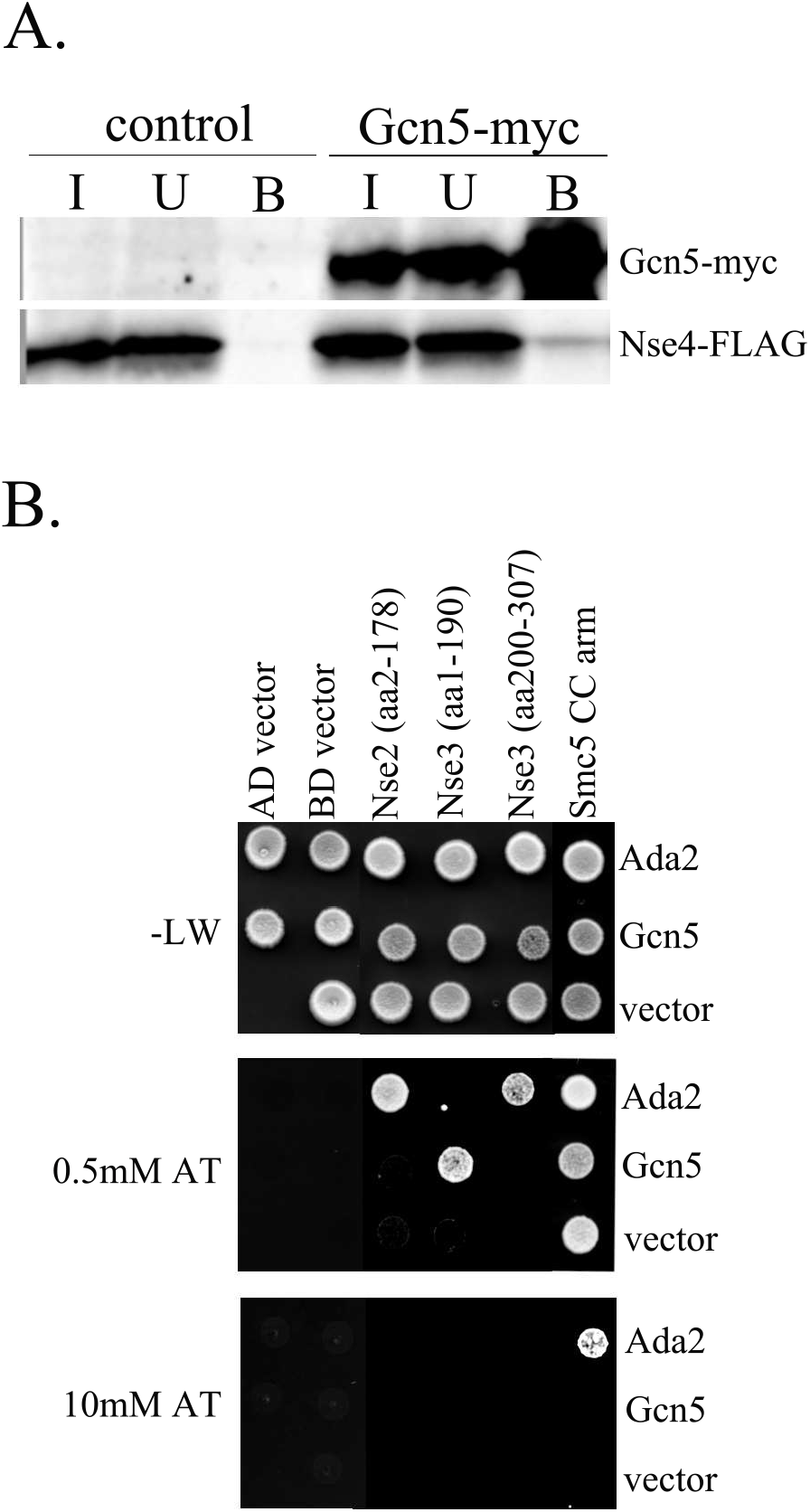
Interactions between the SMC5/6 and SAGA complexes. (**A**) Extracts from fission yeast strains MMP21 (Nse4-FLAG) and YLJ507 (Nse4-FLAG and Gcn5-myc) were immunoprecipitated using the anti-myc antibody. The input (I), unbound (U) and bound (B) fractions were separated by 12% SDS-PAGE. The Nse4-FLAG and Gcn5-myc proteins were analysed on a western blot using anti-FLAG-HRP and anti-myc-HRP, respectively. (**B**) The yeast two-hybrid (Y2H) system was used to determine individual protein-protein interactions between SMC5/6 and SAGA HAT module subunits. The Gal4AD or Gal4BD domains fused to the full-length Ada2 or Gcn5 subunits were co-transformed together with the fragments of SMC5/6 subunits into the PJ69 cells and grown on the plates without Leu, Trp (-L, W; control plates). The protein-protein interactions between SMC5/6 and SAGA were scored by the growth of the yeast PJ69 transformants on the plates without Leu, Trp, and His, containing 3-Amino-1,2,4-triazole (0.5 mM AT or 10 mM AT plates). The fragments were as follows: Gal4BD-Nse2 (aa2-178), Gal4AD-Nse3 (aa1-190), Gal4AD-Nse3 (aa200-307), and Gal4BD-SMC5 CC arm (aa170-225 + 837-910). In control experiments, respective empty pGADT7 (AD) or pGBKT7 (BD) vector was co-transformed with either SAGA or SMC5/6 construct. Note that the Gal4BD-SMC5 CC arm construct self-activated (SMC5-vector combination) and was therefore grown on 10 mM AT plates to assess its binding to Ada2 (SMC5-Ada2 combination).

Next, we used our panel of yeast two-hybrid (Y2H) SMC5/6 subunits to test their interactions with the HAT module (Ada2, Ada3, Gcn5, and Sgf29) subunits. Two of the HAT module subunits, Ada2 and Gcn5, bound the Nse3 subunit (Figs. 2B and S2B). The fragment analysis showed that Gcn5 bound the N-terminal part of Nse3(aa1-190), while Ada2 interacted with its C-terminal WHB domain (Nse3(aa200-307); Figs. 2B and S2C; [45]). In addition, Ada2 bound the N-terminal region of Nse2 and the coiled-coil arm of SMC5 next to the SMC5-Nse2 binding interface [46]. Altogether, our data suggest several physical interactions underlying a functional relationship between SAGA and SMC5/6.

### SMC5/6 and SAGA are required for efficient DNA repair

The SAGA complex plays an important role in transcription regulation and DNA repair by facilitating the accessibility of chromatin to transcription factors and repair proteins [42, 47–50]. Deletion mutants of most SAGA genes were sensitive to hydroxyurea (HU; [42]) and SAGA mutations increased the sensitivity of the *smc5/6* hypomorphic mutants to HU and other DNA-damaging agents (Figs. 3A, S2A, and S3A). These data suggest either a direct role of the SAGA complex in facilitating SMC5/6 access to chromatin at sites of DNA damage or its indirect, independent role.

**Figure 3.**
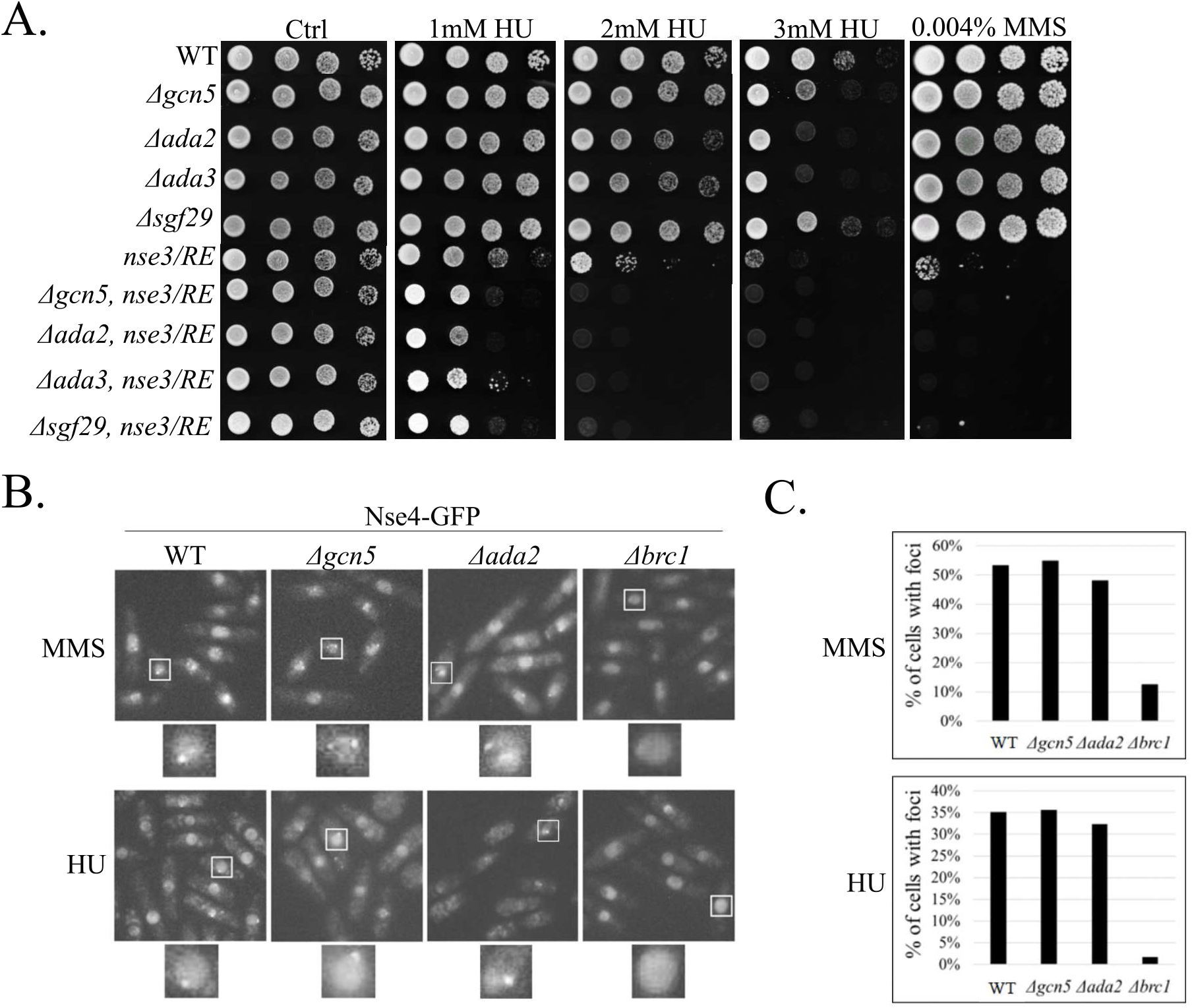
SMC5/6 and SAGA are required for efficient DNA repair. (**A**) Sensitivity of the SMC5/6 and SAGA mutants to genotoxins. Ten-fold serial dilutions of the yeast strains were plated onto YES media containing indicated concentrations of the hydroxyurea (HU) or methyl methane sulfonate (MMS). The double mutants were more sensitive than their respective single mutant counterparts, suggesting the non-redundant functions of SMC5/6 and SAGA in DNA repair. (**B**) Live-cell microscopy of endogenous Nse4-GFP upon HU and MMS treatment, respectively. The Nse4-GFP foci were present in the WT and Δ*gcn5* cells but largely absent in Δ*brc1* cells. (**C**) Quantification of the data in panel B suggests that the localization of SMC5/6 to the DNA-damage foci is independent of SAGA.

Upon DNA damage, SMC5/6 accumulates in foci in a Brc1-dependent way [51, 52]. To assess whether SMC5/6 chromatin accessibility at these DNA damage sites is directly facilitated by SAGA, we analysed the Nse4-GFP foci in the Δ*gcn5* and Δ*ada2* mutants (Fig. 3B). We observed no difference between the frequency of cells with foci in the wild-type (WT), Δ*gcn5*, and Δ*ada2* after treatment with either MMS (methyl methane sulfonate) or HU. In contrast, the number of foci in the Δ*brc1* mutant was strongly reduced (Fig. 3C). These results indicate no direct involvement of SAGA in SMC5/6 localization to sites of DNA damage. Instead, they suggest that the observed HU phenotype results from additive effects of the SAGA and SMC5/6 complexes during DNA damage repair.

### SAGA facilitates chromatin accessibility to SMC5/6 in gene regions

In addition to the accessibility of chromatin to DNA repair and transcription factors, SAGA facilitates chromatin accessibility to condensin complexes [23]. Therefore, we determined the SMC5/6 localization in the unchallenged WT, Δ*gcn5*, Δ*ada2*, and Δ*ubp8* cells using chromatin immunoprecipitation of Nse4-FLAG followed by deep sequencing (ChIP-seq). In the WT cells, most SMC5/6 localized to the repetitive regions (like rDNA, centromeres, or tDNA copies), with the highest occupancy at the rDNA repeats (Fig. 4A) consistent with previous reports [53–58]. Interestingly, we identified 331 Nse4-FLAG peaks (representing ¼ of the total Nse4 occupancy; Fig. 4A) that localized to gene regions.

**Figure 4.**
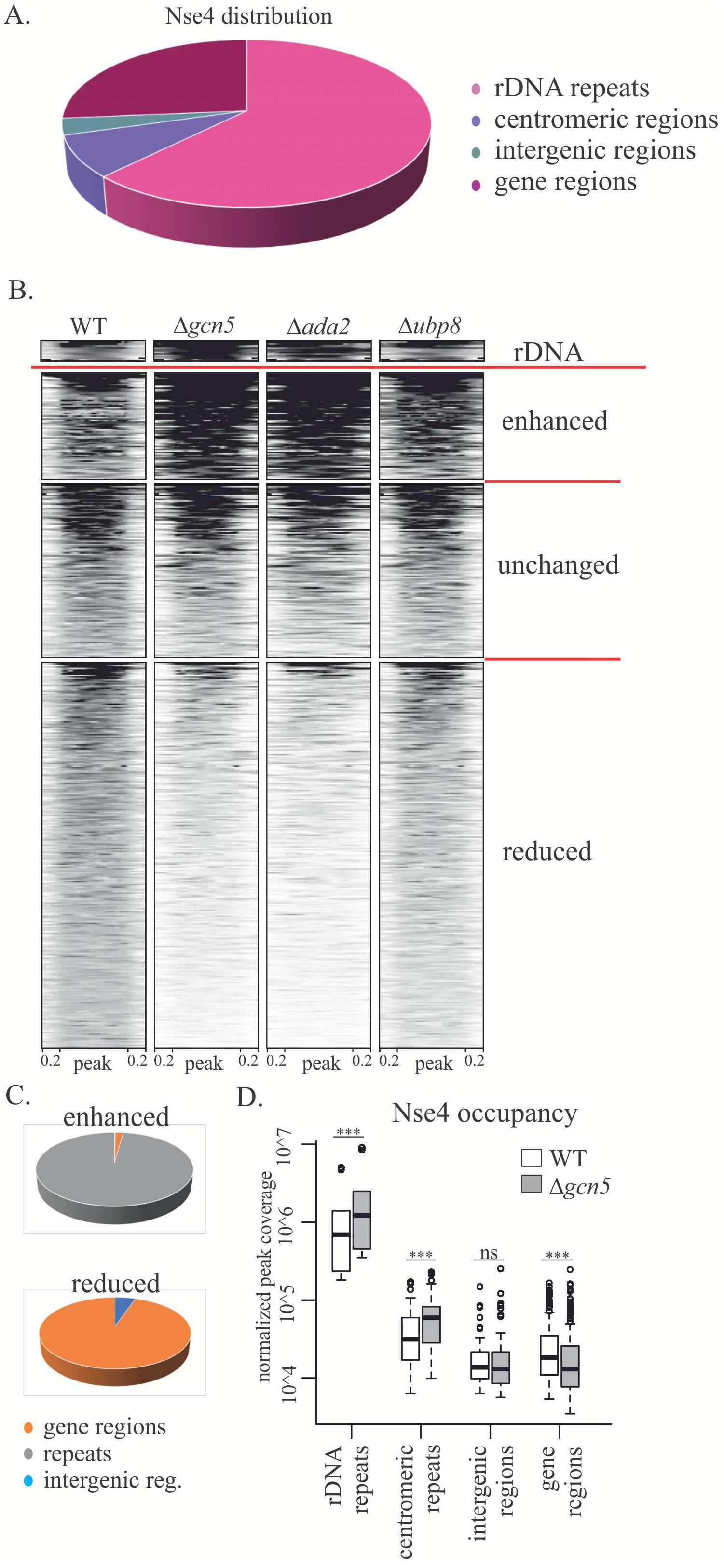
SMC5/6 distribution is dependent on the SAGA HAT module. (**A**) The pie chart shows the distribution of the Nse4-FLAG peak areas in the different genome regions in the WT fission yeast cells. Most SMC5/6 is localized to the repetitive regions (like rDNA and centromeres), with the highest occupancy of the rDNA repeats. A significant portion of the Nse4-FLAG is localized within the intergenic regions or genes. (**B**) The heatmap diagrams compare the occupancy of Nse4-FLAG peaks in the WT, Δ*gcn5*, Δ*ada2*, and Δ*ubp8* mutant cells (as identified in the WT). The top part shows peaks enhanced in Δ*gcn5* and Δ*ada2* (enhanced), while the bottom part clusters peaks reduced in the SAGA HAT module deficient cells (reduced). Peaks in the rDNA repeats are shown separately as these chromosome regions are not fully assembled and annotated in the *S. pombe* reference genome and exert a different range of coverage values (rDNA). The Nse4-FLAG peaks (normalized to median 760 bp width) and their surrounding regions (200 bp upstream and 200 bp downstream) are shown. (**C**) The pie charts show the distribution of the enhanced (top) and reduced (bottom) Nse4-FLAG peak areas in Δ*gcn5*. The SMC5/6 accumulation is mainly enhanced at the repetitive loci. The SMC5/6 localization is primarily reduced in gene regions. (**D**) The box plot graph compares the Nse4-FLAG occupancy in WT and Δ*gcn5* cells. The paired two-sided Wilcoxon statistical test was used: ns, non-significant; ***, p < 0.001.

In Δ*gcn5* and Δ*ada2* mutants, SMC5/6 distribution was altered, while SMC5/6 occupancy in the Δ*ubp8* mutant was similar to that in WT cells. A heatmap clustering analysis showed that the Nse4-FLAG peaks were either enhanced, unchanged, or reduced in the Δ*gcn5* and Δ*ada2* mutants (Fig. 4B). The peaks were mainly enhanced at the repetitive sequences (Fig. 4C), while most peaks in the intergenic regions were not changed (Fig. 4D). Strikingly, most peaks within the gene regions showed reduced Nse4 occupancy (252 out of 331; Figs. 4C and D), suggesting that the SAGA HAT module may facilitate chromatin accessibility to SMC5/6 at gene loci.

The SAGA HAT module acetylates histone H3 at its lysine K9 and K14 residues [59, 60]. To determine the H3K9ac distribution in our fission yeast cells, we performed ChIP-seq using an anti-H3K9ac antibody. A heatmap clustering analysis showed that the magnitude of decrease in SMC5/6 occupancy in gene bodies in the Δ*gcn5* mutant (Δ*gcn5-*WT plot; Figs. 5A and B) correlated with H3K9ac levels around the transcription start site and with transcript levels [61]. It suggests that the SAGA-dependent H3K9 acetylation may facilitate the accessibility of chromatin to SMC5/6 at gene regions.

**Figure 5.**
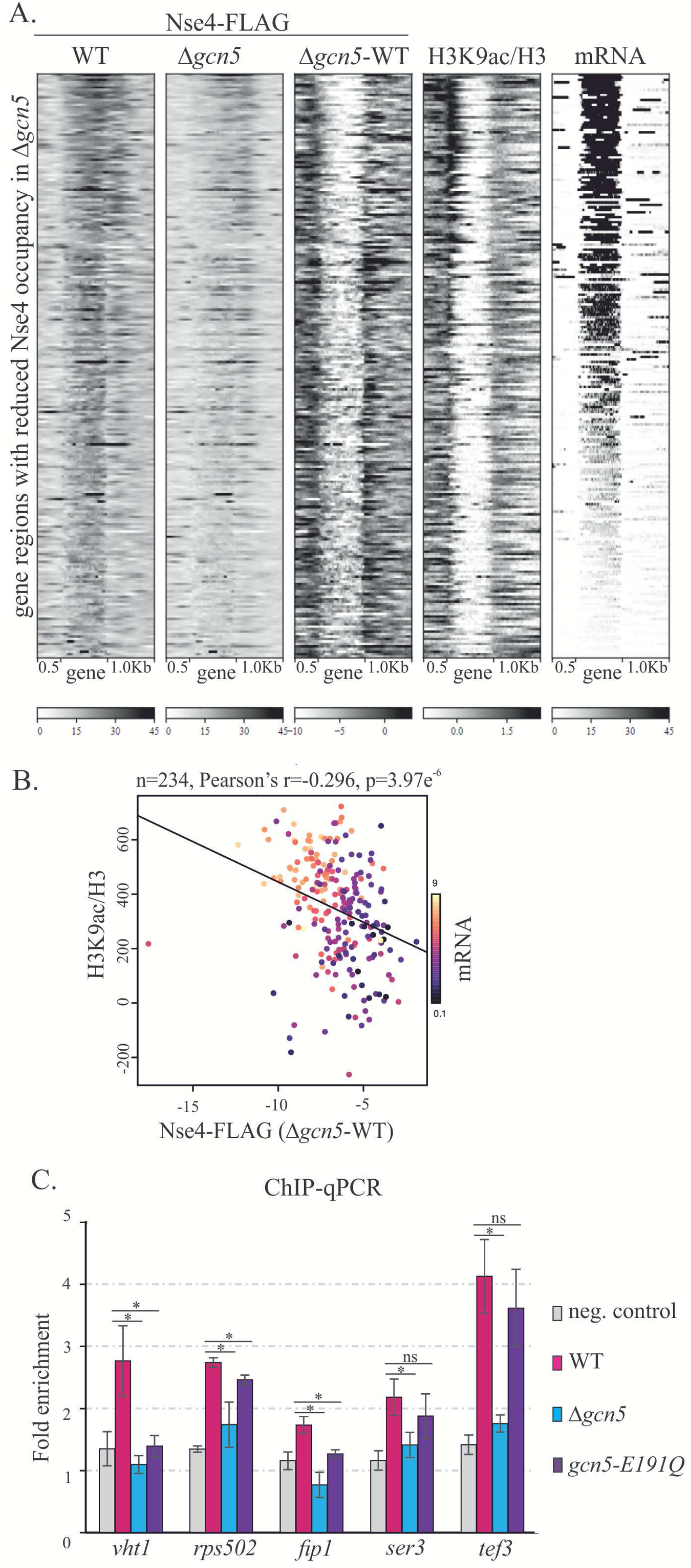
The SMC5/6 accumulation correlates with the H3K9 acetylation status. (**A**) Heatmap statistical analysis of the loci with reduced SMC5/6 occupancy upon the *gcn5* deletion. The Nse4-FLAG signals from WT and Δ*gcn5* are compared, and their differential plot is shown in the middle panel (Δ*gcn5-*WT). Results from ChIP-seq analysis of H3 and H3K9ac are shown (H3K9ac/H3). In addition, transcriptomic (RNA-seq) data are included (mRNA). The genes (normalized to 1 kb width) and their surrounding regions (500bp upstream of TSS and 1 kb downstream of TTS) are shown. (**B**) Scatter plot analysis shows a strong correlation between the drop of SMC5/6 accumulation upon *gcn5* deletion (X-axis) and H3K9-acetylation status (Y-axis) of gene regions (and their transcription levels; colour scale). (**C**) Results of chromatin immunoprecipitation followed by quantitative PCR (ChIP–qPCR) at selected gene loci are shown. Strains 503 (neg. control), MMP21 (WT, containing Nse4-FLAG), Nse4-FLAG Δ*gcn5* (Δ*gcn5*), and Nse4-FLAG *gcn5-E191Q (gcn5-E191Q*) were analysed. The fold enrichment was calculated against the negative *slx9* locus (mean ± standard deviation, n ≥3 biological replicates). The unpaired Wilcoxon statistical test was used: ns, non-significant; *, p < 0.05.

To assess the role of the SAGA-dependent acetylation further, we used the *gcn5*-*E191Q* acetyltransferase-dead mutant [50]. First, we found that the *gcn5*-*E191Q ts* phenotype was exacerbated by *nse3-R254E*, similar to Δ*gcn5* (Suppl. Fig. S3D). Second, the *gcn5*-*E191Q nse3-R254E* double mutant was as sensitive to DNA-damaging agents as the Δ*gcn5 nse3-R254E* double mutant. It suggests that the Gcn5 acetyltransferase activity plays a crucial role in the SAGA-SMC5/6 genetic interactions. Finally, using ChIP-qPCR analysis, we observed reduced Nse4-FLAG occupancy in the *gcn5*-*E191Q* mutant at selected gene regions similar to Δ*gcn5* (Fig. 5C), although some loci exhibited only a modest drop in SMC5/6 levels. Altogether, our results suggest an important role for the SAGA HAT module in facilitating the accessibility of chromatin to SMC5/6 at gene loci.

## Discussion

Here, we discovered the SAGA HAT module as a new chromatin factor assisting in the localization of SMC5/6 to specific genomic loci. In line with SAGA’s best-known function in transcription [5], Δ*gcn5* deletion reduced SMC5/6 occupancy specifically at the gene loci (Fig. 4). In addition, the magnitude of decrease in SMC5/6 occupancy in gene bodies correlated with the H3K9 acetylation and transcript levels (Fig. 5). These data suggest that the SAGA-dependent H3K9 acetylation may facilitate the accessibility of chromatin to SMC5/6. Consistent with this notion, SMC5/6 occupancy was also reduced in the *gcn5*-*E191Q* acetyltransferase-dead mutant. The fact that occupancy reduction was less pronounced in the *gcn5*-*E191Q* mutant could be explained by the physical interactions between SAGA and SMC5/6. Such interactions may target SMC5/6 to SAGA-rich regions, while H3K9 acetylation opening chromatin would enable DNA entrapment. In the *gcn5*-*E191Q* mutant, SMC5/6 could still be targeted to gene regions, although (unmodified) inaccessible chromatin would hamper DNA entrapment. As the Nse3-mediated binding to free DNA is essential for the SMC5/6 function, the DNA-unbound SMC5/6 in the *gcn5*-*E191Q* mutant would likely cause a similar defect as in the Δ*gcn5* mutant and result in the same phenotypes of *gcn5*-*E191Q nse3-R254E* and Δ*gcn5 nse3-R254E* double mutants (Suppl. Fig. S3D and E; [16, 21]).

The additive phenotypes of the double mutants (Fig. 3) probably stem from the non-redundant functions of SMC5/6 and SAGA complexes. For example, SMC5/6 is essential for replication processes (like replication restart; [34]), while SAGA is necessary for the timely transcription of S-phase specific factors [59]. Defects in these functions probably combine in double mutants to cause high HU sensitivity (Figs. 3, S3A, and D). Similarly, the reduced DNA-binding affinity of the *nse3-R254E* mutant synergizes with the reduced binding to gene regions upon *gcn5* deletion, resulting in the slow growth of double mutants. Interestingly, enhanced SMC5/6 accumulation at repetitive regions (Fig. 4) could not rescue these growth defects suggesting an important role of SMC5/6 binding to the gene regions.

It was recently shown that plant ADA2b binds SMC5 and assists in the localization of SMC5/6 to DNA-damage sites [62]. Later studies uncovered an ADA2b-diRNA specific pathway targeting SMC5/6 to DNA-damage foci in plants [63]. However, whether this pathway involves other SAGA components and acetyltransferase activity is unclear. In fission yeast, we and others showed that the SMC5/6 targeting to DNA-damage sites depends on the BRCT domain-containing Brc1 protein but not on Ada2 or Gcn5 (Fig. 3; [64]), suggesting a plant-specific function of the ADA2b-SMC5 interaction in DNA-damage response. Future studies may show if SAGA likewise facilitates the accessibility of chromatin to SMC5/6 at gene loci in plants. Interestingly, we found an interaction between human hADA2b and hNSE4a/b subunits (Suppl. Fig. S2D), suggesting physical interaction between the SAGA and SMC5/6 complexes is conserved in humans. It will be interesting to examine the functional relationships between human SAGA and SMC5/6 complexes, particularly in facilitating SMC5/6 targeting to gene regions.

Consistent with the lack of effect of SAGA on SMC5/6 localization to sites of DNA damage, the SMC5/6 levels at repetitive regions, which are prone to DNA damage even without any genotoxic treatment, were not reduced in Δ*gcn5* and Δ*ada2* deletion mutants (Fig. 4). Furthermore, the SMC5/6 accumulation at repetitive regions depends on H3K9 methyltransferase Clr4 in fission yeast [53]. In our ChIP-seq experiments, the repetitive loci were enriched for SMC5/6 upon SAGA HAT deletion (Fig. 4). We speculate that reduced acetylation levels promoted an increase in methylation levels, indirectly stimulating the SMC5/6 localization to repetitive heterochromatic regions [53, 65]. Altogether, our findings support previous views that different factors target SMC5/6 to different genomic regions [34, 52, 53] and show a new role of SAGA in SMC5/6 targeting to gene regions.

## Materials and methods

### Yeast techniques

Standard fission yeast genetic techniques were used [66]. Yeast strains were crossed and sporulated either at 25°C (*ts* mutants) or 28°C (non-*ts* mutants). Tetrad analysis was carried out on Singer MSM300 (Singer, UK). The deletion integrations were verified on both ends by PCR with specific primers to the G418 cassette and genomic sequence of a deleted gene (approximately 600-800bp from start or end). The PCR products were sequenced.

*S. pombe* cultures were grown to the mid-log phase, and serial 10-fold dilutions were spotted onto rich media with the indicated dose of DNA-damaging agent (hydroxyurea or methyl methane sulfonate). Subsequently, plates were incubated at the indicated temperatures (25, 28, or 37°C) for 3–4 days. Selective media were supplemented with Nourseothricin (cloNAT, 100 μg/ml, Jena Bioscience), G418 (100 μg/ml; Applichem), and/or cycloheximide (100 μg/ml; Sigma).

### Yeast genetic screens

The pAW8-Nse3 integration construct [21] was modified for use in the PEM2 strain as follows. The *Sph*I site was mutated to *Xho*I using a site-directed mutagenesis kit (Agilent Technologies; primers: LJ48 and LJ49; Suppl. Table ST4). The cloNAT cassette was amplified (LJ42 and LJ43) and inserted into the *Xho*I site in front of the Nse3 gene using the In-fusion cloning kit (Takara). A 650bp-long genomic sequence (upstream of Nse3; LJ44 and LJ45) was inserted in front of the cloNAT cassette (using *Xho*I) to ensure its proper integration into the *S. pombe* genome. The mutant cloNAT-*nse3-R254E* construct was created using site-directed mutagenesis (R254E_F and R254E_R primers; [21]). For the yeast transformation, the WT and mutant cloNAT constructs were cleaved by *Spe*I, and the 3246bp long fragment was purified from agarose gel by Gel Extraction Kit (Qiagen). Approximately 1 μg of purified DNA was transformed into the PEM2 strain [30] by standard LiAc protocol. The proper integration of the cloNAT-Nse3 cassette and *rlp42* mutation was checked by PCR and sequencing. The *nse3*-*R254E* PEM2 strain phenotypes were compared with the original *nse3*-*R254E* strain (Suppl. Fig. S1A; [21])

The WT and *nse3*-*R254E* mutant PEM2 strains (YLJ222 and YLJ228; Suppl. Table ST5) were crossed with the *S. pombe* haploid deletion library (BIONEER, version 5, us.bioneer.com) according to the published protocol [30]. The screen was repeated twice using the Rotor HDA robot (Singer, UK). The plate images were taken by a Canon EOS Rebel T3i camera, and the individual colony size was measured. The viability of single deletion mutants (control WT plates) against double mutants (test *nse3*-*R254E* plates) was compared using SGAtools online platform [32]. Genes with a score less than −0.25 were chosen as potential negative interactors (Suppl. Tab. ST1).

The resulting group of 79 genes was analysed by the Gene ontology tool BiNGo [67], which is a plugin of the Cytoscape online platform [33]. The genes were classified according to the Pombase GO database into Biological processes and Cellular component categories, respectively [68]. The default parameters with a 0.05 significance level were applied for both categories.

### Yeast two-hybrid analysis

The Gal4-based Y2H system was used to analyse SMC5/6-SAGA interactions [69]. *S. pombe ada2*, *ada3*, *gcn5*, and *sgf29* genes were PCR amplified from genomic DNA (primers used for *ada2* and *gcn5* cloning are listed in Suppl. Table ST4). All inserts were cloned into respective sites of the pGBKT7 or pGADT7 vectors using the In-Fusion cloning system. pGBKT7-Nse2(aa2-178) was described in [8]. pGADT7-Nse3(aa1-190) was prepared by mutagenesis of 191^st^ aa to STOP codon in pGADT7-Nse3(aa1-328) [70]. The Nse3(aa200-307) fragment was cut out from pTriEx4-Nse3(aa200-307) [45] by *Nco*I-*Xho*I enzymes and cloned into pGADT7. The Smc5(aa170-225+837-910) fragment was amplified from the Smc5(aa2-225+837-1065) construct [71] and inserted into the *Nco*I-*Not*I sites of pGBKT7.

The pairs of pGBKT7 and pGADT7 constructs were co-transformed into the *Saccharomyces cerevisiae* PJ69–4a strain by standard LiAc transformation protocol and selected on SD -Leu, -Trp plates. Drop tests were carried out on SD -Leu, -Trp, -His (with 0.3, 0.5, 1, 3, 5, or 10 mM 3-aminotriazole) plates at 28°C. Each combination of partners was co-transformed and tested at least twice.

### Co-immunoprecipitation of *S. pombe* proteins

Logarithmically growing YLJ507 and MMP21 cells (Suppl. Table ST5; Suppl. Fig. S3B) were cultivated in a rich medium at 28°C (OD_595_ = 0.4 – 0.7). 5×10^8^ cells were harvested by centrifugation (3 min, 4°C, 5000 rpm) and washed with 10 ml of ice-cold PBS. Pellets were stored in the 2 ml screw cup tubes at −80°C. The crude yeast extracts were prepared in 400 μl CHIP lysis buffer (50 mM HEPES, pH 7.5, 140 mM NaCl, 1 mM EDTA, 1% Triton X-100, Complete EDTA-free protease inhibitor cocktail tablets, Roche) with half volume of glass beads (Sigma) in 2 ml low binding tubes using FASTprep-24 (MP Biomedicals; 5 times, 30 sec, 6.5 speed). The suspension was recovered by piercing the bottom of the tube with a needle, placing it into a new 2 ml tube, and centrifugation (3 min, 4°C, 5000 rpm). The beads were washed with 200 μl of CHIP lysis buffer (3 min, 4°C, 5000 rpm). The collected suspensions were clarified by centrifugation (15 min, 4°C, 15000 rpm), and the supernatant was transferred to new low-binding tubes. 40 μl of cell extract was taken for input control. Immunoprecipitation was carried out by adding 2 μl mouse anti-myc antibody (2276S, Cell Signalling) to the cell extracts and incubation for 2 hours at 4°C. 20 μl of protein G coated Dynabeads (Invitrogen) were washed twice with 1 ml CHIP lysis buffer and resuspended in 60 μl of CHIP lysis buffer, then added to the extract with anti-myc antibody and incubated overnight at 4°C. The beads were pelleted using a magnetic rack, and the unbound fraction (40 μl) was taken. Beads were washed four times with 1 ml CHIP lysis buffer, and proteins were eluted by 40 μl of 1xSDS loading buffer. After 15 min incubation at room temperature, the supernatant (bound fraction) was recovered. All fractions were analysed by western blotting using mouse anti-myc-HRP (R951-25, Thermo Fisher) and mouse anti-FLAG-HRP (F1804-1MG, Sigma) antibodies, respectively.

### Chromatin immunoprecipitation analysis (ChIP)

#### Nse4-FLAG ChIP-seq

All strains (Suppl. Table ST5) were cultivated into the mid-log phase (OD = 0.4 – 0.6) and incubated with 1 % formaldehyde for 15 min at room temperature to cross-link DNA–protein complexes. Glycine was added to a final concentration of 125 mM, and the incubation continued for 5 min. 5×10^8^ cells were harvested and washed with 10 ml of ice-cold PBS. The yeast cell wall breakage was performed in 400 μl CHIP lysis buffer with half the volume of glass beads in 2 ml low binding tubes using FASTprep-24. The suspension was washed two times with CHIP lysis buffer (15 min, 4°C, 15000 rpm), and 300 μl of the extract was sonicated with Bioruptor (Diagenode, 30 sec ON/30 sec OFF, High Power, 25 times) and clarified by centrifugation (15 min, 4°C, 15000 rpm), resulting in an average DNA fragment size of 300-500 bp. 5 μl of the sonicated precleared extract was taken as an input control sample.

Monoclonal anti-FLAG M2 antibody (F1804, Sigma) was diluted 1:150, incubated with precleared cell extract in 1.5 ml low binding tube for 2 h on ice, and precipitated overnight with Dynabeads protein G (Invitrogen). Precipitates were washed with 1 ml of CHIP lysis buffer, 1 ml of High Salt buffer (CHIP lysis buffer with 500 mM NaCl), 1 ml of Wash buffer (10 mM Tris-HCl at pH 8.0, 0.25 M LiCl, 0.5% NP-40, 1 mM EDTA), and 1 ml of TE buffer (20 mM Tris-HCl at pH 8.0, 1 mM EDTA). After elution (50 mM Tris at pH 8, 0.1% SDS, 10 mM EDTA) and de-crosslinking overnight at 65°C, the DNA was purified using QIAquick PCR Purification Kit (Qiagen).

For ChIP-seq analysis, the input DNA samples were tested for DNA fragmentation and determination of DNA concentration by the Fragment analyser (Agilent). Input and immunoprecipitation (IP) samples with the best fragmentation and high concentration were used for the creation of NGS libraries (NEBNEXT ULTRA II DNA Library Prep kit, NEB) and sequencing (Illumina Next seq 500, Illumina).

#### H3K9ac/H3 ChIP-seq

Two independent replicates were performed. Cells were grown to the exponential phase (OD_600_ = 0.5) in the complex YES medium and fixed by adding formaldehyde to the final concentration of 1 %. After 30 min incubation, the remaining formaldehyde was quenched by 125 mM glycine. Cells were washed with PBS and broken with glass beads. Extracted chromatin was sheared with the Bioruptor sonicator (Diagenode) using 15 or 30 cycles (for biological replicate 1 and 2, respectively) of 30 sec ON/30 sec OFF at high power settings. For all immunoprecipitations (IP) within a biological replicate, the same amount of chromatin extract was used (2.5 or 3.7 mg of total protein); 1/10 of the total chromatin extract amount was kept for input DNA control. For each IP, 5 μg of antibody (H3: Ab1791, H3K9ac: Ab4441, all Abcam) were incubated with the chromatin extract for 1 hour at 4°C with rotation. Then, 50 μl of BSA-blocked Protein A-coated magnetic beads (10002D, ThermoFisherScientific) were added to the chromatin extract-antibody suspension and incubated for additional 4 hours at 4°C with rotation. The precipitated material and input chromatin extract were de-crosslinked and treated with RNase A and proteinase K. DNA was purified using phenol-chloroform extraction and sodium acetate/ethanol precipitation. In biological replicate 2, DNA purification on AMPure XP beads (AC63880, Beckman Coulter) was performed after the phenol-chloroform extraction to remove low-molecular fragments and RNA. DNA concentration was measured using the Quantus fluorometer (Promega), and fragment size distribution was checked on Agilent Bioanalyser using the High Sensitivity DNA Assay. Library construction and sequencing (50 nt SE) were performed by BGI Tech Solutions (Hong Kong) using the BGISEQ-500 sequencing system.

#### NGS data analysis

The reference fission yeast *S. pombe* genome (2018-09-04) and annotation (2019-11-15) were downloaded from PomBase (https://www.pombase.org/; [68, 72]). Read quality was checked using FastQC version 0.11.8 (https://www.bioinformatics.babraham.ac.uk/projects/fastqc/), and reads were aligned to the *S. pombe* genome using HISAT2 2.1.0 [73] and SAMtools 1.9 [74, 75]. Read coverage tracks (i.e., target protein occupancy) were then computed and normalized to the respective mapped library sizes using deepTools 3.5.1 [76]. The raw ChIP-seq data are available from the ArrayExpress (https://www.ebi.ac.uk/) database under the accession number E-MTAB-11081.

WT fission yeast RNA-seq data were obtained from the NCBI Sequence Read Archive (https://www.ncbi.nlm.nih.gov/sra;datasetsSRR8742773-SRR8742775; [61]). Reads were processed and analysed with the same tools as above. All relevant scripts for (ChIP-seq and RNA-seq) data processing and analysis are available from https://github.com/mprevorovsky/Palecek-Nse-SAGA.

#### ChIP-qPCR

The Nse4-FLAG strains (crossed with the Δ*gcn5* or *gcn5-E191Q* strain; Suppl. Table ST5) were used. The untagged wild-type strain was used as a negative control. All cells were cultivated into the mid-log phase. Cells were then incubated with 1 % formaldehyde for 10 min at room temperature to cross-link DNA–protein complexes. Chromatin immunoprecipitation was performed using a protocol described above for H3K9ac ChIP with the following modifications. Monoclonal anti-FLAG M2 antibody (F1804; Sigma) was diluted at 1:350 (5 μg /sample), incubated with 2 mg of total cell extract for 2 h at 4°C with rotation, and precipitated with Dynabeads protein G (Invitrogen). After overnight incubation, several washes, elution, and de-crosslinking, the DNA was purified using Phenol/Chloroform method.

The relative amount of PCR product was quantified by qPCR using SensiFASTTM SYBR® Hi -ROX Kit (Bioline). The sequences of primers used for the quantitative detection of the chromosomal loci are listed in Suppl Table ST6. Input DNA recovery was calculated as 2 squared [CT(input)− CT(immunoprecipitate)] and normalized to a negative locus *slx9*. Melt curve analysis was performed for each sample after PCR amplification to ensure that a single product was obtained.

### Microscopy

For the Nse4-GFP foci number determination, cells were grown in YES medium overnight, diluted to OD = 0.4 in the morning, and treated with 0.03 % MMS or 20 mM HU for 5 h at 30 °C. 2.5 μl of cell culture was mounted on the slides, and GFP fluorescence was observed. Pictures were taken on the Axio Imager Z1 microscope, using a Plan-Apochromat 63x oil objective, the Axiocam CCD camera, and processed with the AxioVision software (all by Zeiss). A minimum of 500 cells were counted in three independent experiments. For statistical evaluation, p-values were calculated using the χ2 test.

## Acknowledgements

We thank N. J. Krogan and M. N. Boddy for providing the yeast strains. We are grateful to C. Haering for his help with the Rotor HDA robot and M. Adamus for the critical reading of our manuscript.

## Author contributions

Conceptualization: J.J.P., M.P., and D.H.; Data Curation, Formal Analysis, and Funding Acquisition, Supervision: J.J.P. and MP; Investigation: L.M., BŠ., A.M., J.P., P.K., and E.L.; Visualization: J.J.P., M.P. B.Š., and L.M.; Writing – Original Draft Preparation: J.J.P., M.P., P.K., and L.M.

## References

1. Uhlmann F. SMC complexes: from DNA to chromosomes. Nat Rev Mol Cell Biol. 2016;17(7):399–412. Epub 2016/04/14. doi: 10.1038/nrm.2016.30. PubMed PMID: 27075410.

2. Davidson IF, Peters JM. Genome folding through loop extrusion by SMC complexes. Nat Rev Mol Cell Biol. 2021;22(7):445–64. Epub 20210325. doi: 10.1038/s41580-021-00349-7. PubMed PMID: 33767413.

3. Ransom M, Dennehey BK, Tyler JK. Chaperoning histones during DNA replication and repair. Cell. 2010;140(2):183–95. doi: 10.1016/j.cell.2010.01.004. PubMed PMID: 20141833; PubMed Central PMCID: PMCPMC3433953.

4. Morrison AJ, Shen XT. Chromatin remodelling beyond transcription: the INO80 and SWR1 complexes. Nature Reviews Molecular Cell Biology. 2009;10(6):373–84. doi: 10.1038/nrm2693. PubMed PMID: WOS:000266270900012.

5. Helmlinger D, Tora L. Sharing the SAGA. Trends Biochem Sci. 2017;42(11):850–61. Epub 2017/09/27. doi: 10.1016/j.tibs.2017.09.001. PubMed PMID: 28964624; PubMed Central PMCID: PMCPMC5660625.

6. Palecek JJ, Gruber S. Kite Proteins: a Superfamily of SMC/Kleisin Partners Conserved Across Bacteria, Archaea, and Eukaryotes. Structure. 2015;23(12):2183–90. doi: 10.1016/j.str.2015.10.004. PubMed PMID: WOS:000366171500001.

7. Wells JN, Gligoris TG, Nasmyth KA, Marsh JA. Evolution of condensin and cohesin complexes driven by replacement of Kite by Hawk proteins. Curr Biol. 2017;27(1):R17–R8. doi: 10.1016/j.cub.2016.11.050. PubMed PMID: 28073014; PubMed Central PMCID: PMCPMC5228436.

8. Andrews E, Palecek J, Sergeant J, Taylor E, Lehmann A, Watts F. Nse2, a component of the Smc5-6 complex, is a SUMO ligase required for the response to DNA damage. Mol Cell Biol. 2005;25(1):185–96. doi: 10.1128/MCB.25.1.185-196.2005. PubMed PMID: WOS:000226236900016.

9. Zhao X, Blobel G. A SUMO ligase is part of a nuclear multiprotein complex that affects DNA repair and chromosomal organization. Proc Natl Acad Sci U S A. 2005;102(13):4777–47782. PubMed PMID: 15738391.

10. Gligoris T, Löwe J. Structural Insights into Ring Formation of Cohesin and Related Smc Complexes. Trends Cell Biol. 2016;26(9):680–93. Epub 2016/04/28. doi: 10.1016/j.tcb.2016.04.002. PubMed PMID: 27134029; PubMed Central PMCID: PMCPMC4989898.

11. Hassler M, Shaltiel IA, Haering CH. Towards a Unified Model of SMC Complex Function. Curr Biol. 2018;28(21):R1266–R81. doi: 10.1016/j.cub.2018.08.034. PubMed PMID: 30399354.

12. Nasmyth K, Haering CH. The structure and function of SMC and kleisin complexes. Annu Rev Biochem. 2005;74:595–648.

13. Alt A, Dang HQ, Wells OS, Polo LM, Smith MA, McGregor GA, et al. Specialized interfaces of Smc5/6 control hinge stability and DNA association. Nat Commun. 2017;8:14011. Epub 2017/01/30. doi: 10.1038/ncomms14011. PubMed PMID: 28134253; PubMed Central PMCID: PMCPMC5290277.

14. Hallett ST, Campbell Harry I, Schellenberger P, Zhou L, Cronin NB, Baxter J, et al. Cryo-EM structure of the Smc5/6 holo-complex. Nucleic Acids Res. 2022. Epub 20220822. doi: 10.1093/nar/gkac692. PubMed PMID: 35993814; PubMed Central PMCID: PMCPMC9458440.

15. Adamus M, Lelkes E, Potesil D, Ganji SR, Kolesar P, Zabrady K, et al. Molecular Insights into the Architecture of the Human SMC5/6 Complex. Journal of Molecular Biology. 2020;432(13):3820–37. doi: 10.1016/j.jmb.2020.04.024. PubMed PMID: WOS:000541931500007.

16. Yu Y, Li S, Ser Z, Kuang H, Than T, Guan D, et al. Cryo-EM structure of DNA-bound Smc5/6 reveals DNA clamping enabled by multi-subunit conformational changes. Proc Natl Acad Sci U S A. 2022;119(23):e2202799119. Epub 20220601. doi: 10.1073/pnas.2202799119. PubMed PMID: 35648833.

17. Ganji M, Shaltiel IA, Bisht S, Kim E, Kalichava A, Haering CH, et al. Real-time imaging of DNA loop extrusion by condensin. Science. 2018;360(6384):102–5. Epub 2018/02/22. doi: 10.1126/science.aar7831. PubMed PMID: 29472443.

18. Davidson IF, Bauer B, Goetz D, Tang W, Wutz G, Peters JM. DNA loop extrusion by human cohesin. Science. 2019;366(6471):1338–45. Epub 2019/11/21. doi: 10.1126/science.aaz3418. PubMed PMID: 31753851.

19. Wang XD, Hughes AC, Brandao HB, Walker B, Lierz C, Cochran JC, et al. In Vivo Evidence for ATPase-Dependent DNA Translocation by the Bacillus subtilis SMC Condensin Complex. Molecular Cell. 2018;71(5):841–7. doi: 10.1016/j.molcel.2018.07.006. PubMed PMID: WOS:000443829100016.

20. Pradhan B, Kanno K, Igarashi MU, Baaske MD, Wong JSK, Jeppsson K, et al. The Smc5/6 complex is a DNA loop extruding motor. bioRxiv. 2022. doi: 10.1101/2022.05.13.491800.

21. Zabrady K, Adamus M, Vondrova L, Liao C, Skoupilova H, Novakova M, et al. Chromatin association of the SMC5/6 complex is dependent on binding of its NSE3 subunit to DNA. Nucleic Acids Research. 2016;44(3):1064–79. doi: 10.1093/nar/gkv1021. PubMed PMID: WOS:000371268700017.

22. Piazza I, Rutkowska A, Ori A, Walczak M, Metz J, Pelechano V, et al. Association of condensin with chromosomes depends on DNA binding by its HEAT-repeat subunits. Nat Struct Mol Biol. 2014;21(6):560–8. Epub 20140518. doi: 10.1038/nsmb.2831. PubMed PMID: 24837193.

23. Toselli-Mollereau E, Robellet X, Fauque L, Lemaire S, Schiklenk C, Klein C, et al. Nucleosome eviction in mitosis assists condensin loading and chromosome condensation. EMBO J. 2016;35(14):1565–81. Epub 2016/06/06. doi: 10.15252/embj.201592849. PubMed PMID: 27266525; PubMed Central PMCID: PMCPMC4946138.

24. Shaltiel IA, Datta S, Lecomte L, Hassler M, Kschonsak M, Bravo S, et al. A hold-and-feed mechanism drives directional DNA loop extrusion by condensin. Science. 2022;376(6597):1087–94. Epub 20220602. doi: 10.1126/science.abm4012. PubMed PMID: 35653469.

25. Shi ZB, Gao HS, Bai XC, Yu HT. Cryo-EM structure of the human cohesin-NIPBL-DNA complex. Science. 2020;368(6498):1454–+. doi: 10.1126/science.abb0981. PubMed PMID: WOS:000545264600036.

26. Bürmann F, Funke LFH, Chin JW, Löwe J. Cryo-EM structure of MukBEF reveals DNA loop entrapment at chromosomal unloading sites. Mol Cell. 2021;81(23):4891–906.e8. Epub 20211104. doi: 10.1016/j.molcel.2021.10.011. PubMed PMID: 34739874; PubMed Central PMCID: PMCPMC8669397.

27. Lopez-Serra L, Kelly G, Patel H, Stewart A, Uhlmann F. The Scc2-Scc4 complex acts in sister chromatid cohesion and transcriptional regulation by maintaining nucleosome-free regions. Nature Genetics. 2014;46(10):1147–51. doi: 10.1038/ng.3080. PubMed PMID: WOS:000342554100021.

28. Munoz S, Minamino M, Casas-Delucchi CS, Patel H, Uhlmann F. A Role for Chromatin Remodeling in Cohesin Loading onto Chromosomes. Molecular Cell. 2019;74(4):664.+. doi: 10.1016/j.molcel.2019.02.027. PubMed PMID: WOS:000468094300005.

29. Munoz S, Passarelli F, Uhlmann F. Conserved roles of chromatin remodellers in cohesin loading onto chromatin. Current Genetics. 2020;66(5):951–6. doi: 10.1007/s00294-020-01075-x. PubMed PMID: WOS:000558454500001.

30. Roguev A, Wiren M, Weissman JS, Krogan NJ. High-throughput genetic interaction mapping in the fission yeast Schizosaccharomyces pombe. Nat Methods. 2007;4(10):861–6. Epub 2007/09/23. doi: 10.1038/nmeth1098. PubMed PMID: 17893680.

31. Kim DU, Hayles J, Kim D, Wood V, Park HO, Won M, et al. Analysis of a genome-wide set of gene deletions in the fission yeast Schizosaccharomyces pombe. Nat Biotechnol. 2010;28(6):617–23. Epub 2010/05/16. doi: 10.1038/nbt.1628. PubMed PMID: 20473289; PubMed Central PMCID: PMCPMC3962850.

32. Wagih O, Usaj M, Baryshnikova A, VanderSluis B, Kuzmin E, Costanzo M, et al. SGAtools: one-stop analysis and visualization of array-based genetic interaction screens. Nucleic Acids Res. 2013;41(Web Server issue):W591–6. Epub 2013/05/15. doi: 10.1093/nar/gkt400. PubMed PMID: 23677617; PubMed Central PMCID: PMCPMC3692131.

33. Shannon P, Markiel A, Ozier O, Baliga NS, Wang JT, Ramage D, et al. Cytoscape: a software environment for integrated models of biomolecular interaction networks. Genome Res. 2003;13(11):2498–504. doi: 10.1101/gr.1239303. PubMed PMID: 14597658; PubMed Central PMCID: PMCPMC403769.

34. Palecek JJ. SMC5/6: Multifunctional Player in Replication. Genes. 2019;10(1):E7. doi: 10.3390/genes10010007. PubMed PMID: WOS:000459743800007.

35. Aragon L. The Smc5/6 Complex: New and Old Functions of the Enigmatic Long-Distance Relative. In: Bonini NM, editor. Annual Review of Genetics, Vol 52. Annual Review of Genetics. 52. Palo Alto: Annual Reviews; 2018. p. 89–107.

36. Lee KM, Nizza S, Hayes T, Bass KL, Irmisch A, Murray JM, et al. Brc1-mediated rescue of Smc5/6 deficiency: requirement for multiple nucleases and a novel Rad18 function. Genetics. 2007;175(4):1585–95.

37. Lehmann AR, Walicka M, Griffiths DJF, Murray JM, Watts FZ, McCready S, et al. The rad18 gene of Schizosaccharomyces pombe defines a new subgroup of the SMC superfamily involved in DNA repair. Mol Cell Biol. 1995;15:7067–80.

38. Morikawa H, Morishita T, Kawane S, Iwasaki H, Carr AM, Shinagawa H. Rad62 protein functionally and physically associates with the smc5/smc6 protein complex and is required for chromosome integrity and recombination repair in fission yeast. Mol Cell Biol. 2004;24(21):9401–13.

39. Dixon SJ, Fedyshyn Y, Koh JLY, Prasad TSK, Chahwan C, Chua G, et al. Significant conservation of synthetic lethal genetic interaction networks between distantly related eukaryotes. Proceedings of the National Academy of Sciences of the United States of America. 2008;105(43):16653–8. doi: 10.1073/pnas.0806261105. PubMed PMID: WOS:000260913500046.

40. Ryan CJ, Roguev A, Patrick K, Xu J, Jahari H, Tong Z, et al. Hierarchical modularity and the evolution of genetic interactomes across species. Mol Cell. 2012;46(5):691–704. doi: 10.1016/j.molcel.2012.05.028. PubMed PMID: 22681890; PubMed Central PMCID: PMCPMC3380636.

41. Pebernard S, McDonald WH, Pavlova Y, Yates JR, Boddy MN. Nse1, Nse2, and a novel subunit of the Smc5-Smc6 complex, Nse3, play a crucial role in meiosis. Mol Biol Cell. 2004;15(11):4866–76. doi: 10.1091/mbc.E04-05-0436. PubMed PMID: 15331764; PubMed Central PMCID: PMCPMC524734.

42. Helmlinger D, Marguerat S, Villén J, Swaney DL, Gygi SP, Bähler J, et al. Tra1 has specific regulatory roles, rather than global functions, within the SAGA co-activator complex. EMBO J. 2011;30(14):2843–52. Epub 2011/06/03. doi: 10.1038/emboj.2011.181. PubMed PMID: 21642955; PubMed Central PMCID: PMCPMC3160243.

43. Verkade HM, Bugg SJ, Lindsay HD, Carr AM, O’Connell MJ. Rad18 is required for DNA repair and checkpoint responses in fission yeast. Mol Biol Cell. 1999;10(9):2905–18.

44. Kolesar P, Stejskal K, Potesil D, Murray JM, Palecek JJ. Role of Nse1 Subunit of SMC5/6 Complex as a Ubiquitin Ligase. Cells. 2022;11(1):13. doi: 10.3390/cells11010165. PubMed PMID: WOS:000758383300001.

45. Hudson JJR, Bednarova K, Kozakova L, Liao CY, Guerineau M, Colnaghi R, et al. Interactions between the Nse3 and Nse4 Components of the SMC5-6 Complex Identify Evolutionarily Conserved Interactions between MAGE and EID Families. Plos One. 2011;6(2):14. doi: 10.1371/journal.pone.0017270. PubMed PMID: WOS:000287764100039.

46. Duan X, Sarangi P, Liu X, Rangi GK, Zhao X, Ye H. Structural and functional insights into the roles of the Mms21 subunit of the Smc5/6 complex. Mol Cell. 2009;35(5):657–68. PubMed PMID: 19748359.

47. Clouaire T, Rocher V, Lashgari A, Arnould C, Aguirrebengoa M, Biernacka A, et al. Comprehensive Mapping of Histone Modifications at DNA Double-Strand Breaks Deciphers Repair Pathway Chromatin Signatures. Mol Cell. 2018;72(2):250–62.e6. Epub 2018/09/27. doi: 10.1016/j.molcel.2018.08.020. PubMed PMID: 30270107; PubMed Central PMCID: PMCPMC6202423.

48. Deshpande GP, Hayles J, Hoe KL, Kim DU, Park HO, Hartsuiker E. Screening a genome-wide S. pombe deletion library identifies novel genes and pathways involved in genome stability maintenance. DNA Repair (Amst). 2009;8(5):672–9. Epub 2009/03/04. doi: 10.1016/j.dnarep.2009.01.016. PubMed PMID: 19264558; PubMed Central PMCID: PMCPMC2675035.

49. Pan X, Lei B, Zhou N, Feng B, Yao W, Zhao X, et al. Identification of novel genes involved in DNA damage response by screening a genome-wide Schizosaccharomyces pombe deletion library. BMC Genomics. 2012;13:662. Epub 2012/11/23. doi: 10.1186/1471-2164-13-662. PubMed PMID: 23173672; PubMed Central PMCID: PMCPMC3536581.

50. Helmlinger D, Marguerat S, Villén J, Gygi SP, Bähler J, Winston F. The S. pombe SAGA complex controls the switch from proliferation to sexual differentiation through the opposing roles of its subunits Gcn5 and Spt8. Genes Dev. 2008;22(22):3184–95. doi: 10.1101/gad.1719908. PubMed PMID: 19056896; PubMed Central PMCID: PMCPMC2593614.

51. Oravcová M, Gadaleta MC, Nie M, Reubens MC, Limbo O, Russell P, et al. Brc1 Promotes the Focal Accumulation and SUMO Ligase Activity of Smc5-Smc6 During Replication Stress. Mol Cell Biol. 2018. Epub 2018/10/22. doi: 10.1128/MCB.00271-18. PubMed PMID: 30348841.

52. Oravcová M, Boddy MN. Recruitment, loading, and activation of the Smc5-Smc6 SUMO ligase. Curr Genet. 2019;65(3):669–76. Epub 2019/01/02. doi: 10.1007/s00294-018-0922-9. PubMed PMID: 30600397; PubMed Central PMCID: PMCPMC6511331.

53. Pebernard S, Schaffer L, Campbell D, Head SR, Boddy MN. Localization of Smc5/6 to centromeres and telomeres requires heterochromatin and SUMO, respectively. EMBO J. 2008;27(22):3011–23.

54. Lindroos HB, Strom L, Itoh T, Katou Y, Shirahige K, Sjogren C. Chromosomal association of the Smc5/6 complex reveals that it functions in differently regulated pathways. Mol Cell. 2006;22(6):755–67.

55. Irmisch A, Ampatzidou E, Mizuno K, O’Connell MJ, Murray JM. Smc5/6 maintains stalled replication forks in a recombination-competent conformation. EMBO J. 2009;28(2):144–55. Epub 2009/01/23. doi: emboj2008273 [pii] 10.1038/emboj.2008.273. PubMed PMID: 19158664.

56. Peng XP, Lim S, Li SB, Marjavaara L, Chabes A, Zhao XL. Acute Smc5/6 depletion reveals its primary role in rDNA replication by restraining recombination at fork pausing sites. Plos Genetics. 2018;14(1):20. doi: 10.1371/journal.pgen.1007129. PubMed PMID: WOS:000423718600006.

57. Torres-Rosell J, Machin F, Farmer S, Jarmuz A, Eydmann T, Dalgaard JZ, et al. SMC5 and SMC6 genes are required for the segregation of repetitive chromosome regions. Nat Cell Biol. 2005;7(4):412–9.

58. Torres-Rosell J, Sunjevaric I, De Piccoli G, Sacher M, Eckert-Boulet N, Reid R, et al. The Smc5-Smc6 complex and SUMO modification of Rad52 regulates recombinational repair at the ribosomal gene locus. Nat Cell Biol. 2007;9(8):923–31. Epub 2007/07/22. doi: 10.1038/ncb1619. PubMed PMID: 17643116.

59. González-Medina A, Hidalgo E, Ayté J. Gcn5-mediated acetylation at MBF-regulated promoters induces the G1/S transcriptional wave. Nucleic Acids Res. 2019;47(16):8439–51. doi: 10.1093/nar/gkz561. PubMed PMID: 31260531; PubMed Central PMCID: PMCPMC6895280.

60. Nugent RL, Johnsson A, Fleharty B, Gogol M, Xue-Franzén Y, Seidel C, et al. Expression profiling of S. pombe acetyltransferase mutants identifies redundant pathways of gene regulation. BMC Genomics. 2010;11:59. Epub 20100122. doi: 10.1186/1471-2164-11-59. PubMed PMID: 20096118; PubMed Central PMCID: PMCPMC2823694.

61. Elías-Villalobos A, Toullec D, Faux C, Séveno M, Helmlinger D. Chaperone-mediated ordered assembly of the SAGA and NuA4 transcription co-activator complexes in yeast. Nat Commun. 2019;10(1):5237. Epub 2019/11/20. doi: 10.1038/s41467-019-13243-w. PubMed PMID: 31748520; PubMed Central PMCID: PMCPMC6868236.

62. Lai J, Jiang J, Wu Q, Mao N, Han D, Hu H, et al. The transcriptional coactivator ADA2b recruits a structural maintenance protein to double-strand breaks during DNA repair in plants. Plant Physiol. 2018. Epub 2018/02/20. doi: 10.1104/pp.18.00123. PubMed PMID: 29463775.

63. Jiang J, Ou X, Han D, He Z, Liu S, Mao N, et al. A diRNA-protein scaffold module mediates SMC5/6 recruitment in plant DNA repair. Plant Cell. 2022. Epub 20220701. doi: 10.1093/plcell/koac191. PubMed PMID: 35775944.

64. Oravcova M, Gadaleta MC, Nie MH, Reubens MC, Limbo O, Russell P, et al. Brc1 Promotes the Focal Accumulation and SUMO Ligase Activity of Smc5-Smc6 during Replication Stress. Molecular and Cellular Biology. 2019;39(2):15. doi: 10.1128/mcb.00271-18. PubMed PMID: WOS:000454772000001.

65. Chiolo I, Minoda A, Colmenares SU, Polyzos A, Costes SV, Karpen GH. Double-Strand Breaks in Heterochromatin Move Outside of a Dynamic HP1a Domain to Complete Recombinational Repair. Cell. 2011;144(5):732–44. Epub 2011 Feb 25.

66. Moreno S, Klar A, Nurse P. Molecular genetic analysis of fission yeast Schizosaccharomyces pombe. Methods Enzymol. 1991;194:795–823. doi: 10.1016/0076-6879(91)94059-l. PubMed PMID: 2005825.

67. Maere S, Heymans K, Kuiper M. BiNGO: a Cytoscape plugin to assess overrepresentation of gene ontology categories in biological networks. Bioinformatics. 2005;21(16):3448–9. Epub 2005/06/24. doi: 10.1093/bioinformatics/bti551. PubMed PMID: 15972284.

68. Lock A, Rutherford K, Harris MA, Hayles J, Oliver SG, Bähler J, et al. PomBase 2018: user-driven reimplementation of the fission yeast database provides rapid and intuitive access to diverse, interconnected information. Nucleic Acids Res. 2019;47(D1):D821–D7. doi: 10.1093/nar/gky961. PubMed PMID: 30321395; PubMed Central PMCID: PMCPMC6324063.

69. Paleček JJ, Vondrová L, Zábrady K, Otočka J. Multicomponent Yeast Two-Hybrid System: Applications to Study Protein-Protein Interactions in SMC Complexes. Methods Mol Biol. 2019;2004:79–90. doi: 10.1007/978-1-4939-9520-2_7. PubMed PMID: 31147911.

70. Sergeant J, Taylor E, Palecek J, Fousteri M, Andrews E, Sweeney S, et al. Composition and architecture of the Schizosaccharomyces pombe Rad18 (Smc5-6) complex. Mol Cell Biol. 2005;25(1):172–84. doi: 10.1128/MCB.25.1.172-184.2005. PubMed PMID: WOS:000226236900015.

71. Palecek J, Vidot S, Feng M, Doherty AJ, Lehmann AR. The SMC5-6 DNA repair complex: Bridging of the SMC5-6 heads by the Kleisin, NSE4, and non-Kleisin subunits. J Biol Chem. 2006;281:36952–9. PubMed PMID: 17005570.

72. Wood V, Gwilliam R, Rajandream MA, Lyne M, Lyne R, Stewart A, et al. The genome sequence of Schizosaccharomyces pombe. Nature. 2002;415(6874):871–80.

73. Kim D, Langmead B, Salzberg SL. HISAT: a fast spliced aligner with low memory requirements. Nat Methods. 2015;12(4):357–60. Epub 20150309. doi: 10.1038/nmeth.3317. PubMed PMID: 25751142; PubMed Central PMCID: PMCPMC4655817.

74. Li H, Handsaker B, Wysoker A, Fennell T, Ruan J, Homer N, et al. The Sequence Alignment/Map format and SAMtools. Bioinformatics. 2009;25(16):2078–9. Epub 20090608. doi: 10.1093/bioinformatics/btp352. PubMed PMID: 19505943; PubMed Central PMCID: PMCPMC2723002.

75. Bonfield JK, Marshall J, Danecek P, Li H, Ohan V, Whitwham A, et al. HTSlib: C library for reading/writing high-throughput sequencing data. Gigascience. 2021;10(2). doi: 10.1093/gigascience/giab007. PubMed PMID: 33594436; PubMed Central PMCID: PMCPMC7931820.

76. Ramírez F, Ryan DP, Grüning B, Bhardwaj V, Kilpert F, Richter AS, et al. deepTools2: a next generation web server for deep-sequencing data analysis. Nucleic Acids Res. 2016;44(W1):W160–5. Epub 20160413. doi: 10.1093/nar/gkw257. PubMed PMID: 27079975; PubMed Central PMCID: PMCPMC4987876.

77. Tapia-Alveal C, Lin SJ, Yeoh A, Jabado OJ, O’Connell MJ. H2A.Z-Dependent Regulation of Cohesin Dynamics on Chromosome Arms. Mol Cell Biol. 2014;34(11):2092–104.

78. Bonnet J, Wang CY, Baptista T, Vincent SD, Hsiao WC, Stierle M, et al. The SAGA coactivator complex acts on the whole transcribed genome and is required for RNA polymerase II transcription. Genes Dev. 2014;28(18):1999–2012. doi: 10.1101/gad.250225.114. PubMed PMID: 25228644; PubMed Central PMCID: PMCPMC4173158.

79. Bavner A, Matthews J, Sanyal S, Gustafsson JA, Treuter E. EID3 is a novel EID family member and an inhibitor of CBP-dependent co-activation. Nucleic Acids Res. 2005;33(11):3561–9. PubMed PMID: 15987788.

80. Schneider M, Hellerschmied D, Schubert T, Amlacher S, Vinayachandran V, Reja R, et al. The Nuclear Pore-Associated TREX-2 Complex Employs Mediator to Regulate Gene Expression. Cell. 2015;162(5):1016–28. doi: 10.1016/j.cell.2015.07.059. PubMed PMID: 26317468; PubMed Central PMCID: PMCPMC4644235.

81. Luthra R, Kerr SC, Harreman MT, Apponi LH, Fasken MB, Ramineni S, et al. Actively transcribed GAL genes can be physically linked to the nuclear pore by the SAGA chromatin modifying complex. J Biol Chem. 2007;282(5):3042–9. Epub 2006/12/06. doi: 10.1074/jbc.M608741200. PubMed PMID: 17158105.

82. Brown CE, Howe L, Sousa K, Alley SC, Carrozza MJ, Tan S, et al. Recruitment of HAT complexes by direct activator interactions with the ATM-related Tra1 subunit. Science. 2001;292(5525):2333–7. doi: 10.1126/science.1060214. PubMed PMID: 11423663.

83. Rivosecchi J, Jost D, Vachez L, Gautier FDR, Bernard P, Vanoosthuyse V. RNA polymerase backtracking results in the accumulation of fission yeast condensin at active genes. Life Science Alliance. 2021;4(6):12. doi: 10.26508/lsa.202101046. PubMed PMID: WOS:000654748200015.

84. Nakazawa N, Arakawa O, Yanagida M. Condensin locates at transcriptional termination sites in mitosis, possibly releasing mitotic transcripts. Open Biology. 2019;9(10):9. doi: 10.1098/rsob.190125. PubMed PMID: WOS:000521587300003.

85. Hocquet C, Robellet X, Modolo L, Sun XM, Burny C, Cuylen-Haering S, et al. Condensin controls cellular RNA levels through the accurate segregation of chromosomes instead of directly regulating transcription. Elife. 2018;7:31. doi: 10.7554/eLife.38517. PubMed PMID: WOS:000446417500001.

86. Kim KD, Tanizawa H, Iwasaki O, Noma K. Transcription factors mediate condensin recruitment and global chromosomal organization in fission yeast. Nat Genet. 2016;48(10):1242–52. Epub 2016/08/22. doi: 10.1038/ng.3647. PubMed PMID: 27548313; PubMed Central PMCID: PMCPMC5042855.

87. Lengronne A, Katou Y, Mori S, Yokobayashi S, Kelly GP, Itoh T, et al. Cohesin relocation from sites of chromosomal loading to places of convergent transcription. Nature. 2004;430(6999):573–8. Epub 2004 Jun 30.

88. D’Ambrosio C, Schmidt CK, Katou Y, Kelly G, Itoh T, Shirahige K, et al. Identification of cis-acting sites for condensin loading onto budding yeast chromosomes. Genes Dev. 2008;22(16):2215–27. doi: 10.1101/gad.1675708. PubMed PMID: 18708580; PubMed Central PMCID: PMCPMC2518811.

89. Kakui Y, Barrington C, Barry DJ, Gerguri T, Fu X, Bates PA, et al. Fission yeast condensin contributes to interphase chromatin organization and prevents transcription-coupled DNA damage. Genome Biol. 2020;21(1):272. Epub 2020/11/05. doi: 10.1186/s13059-020-02183-0. PubMed PMID: 33153481; PubMed Central PMCID: PMCPMC7643427.

90. Borrie MS, Campor JS, Joshi H, Gartenberg MR. Binding, sliding, and function of cohesin during transcriptional activation. Proc Natl Acad Sci U S A. 2017;114(7):E1062–e71. Epub 20170130. doi: 10.1073/pnas.1617309114. PubMed PMID: 28137853; PubMed Central PMCID: PMCPMC5320966.

91. Kong M, Cutts EE, Pan D, Beuron F, Kaliyappan T, Xue C, et al. Human Condensin I and II Drive Extensive ATP-Dependent Compaction of Nucleosome-Bound DNA. Mol Cell. 2020;79(1):99–114.e9. Epub 20200522. doi: 10.1016/j.molcel.2020.04.026. PubMed PMID: 32445620; PubMed Central PMCID: PMCPMC7335352.

92. Johnsson A, Durand-Dubief M, Xue-Franzén Y, Rönnerblad M, Ekwall K, Wright A. HAT-HDAC interplay modulates global histone H3K14 acetylation in gene-coding regions during stress. EMBO Rep. 2009;10(9):1009–14. Epub 2009/07/24. doi: 10.1038/embor.2009.127. PubMed PMID: 19633696; PubMed Central PMCID: PMCPMC2750064.

93. Garcia-Luis J, Lazar-Stefanita L, Gutierrez-Escribano P, Thierry A, Cournac A, Garcia A, et al. FACT mediates cohesin function on chromatin. Nature Structural & Molecular Biology. 2019;26(10):970–+. doi: 10.1038/s41594-019-0307-x. PubMed PMID: WOS:000488970400021.

94. Shintomi K, Takahashi TS, Hirano T. Reconstitution of mitotic chromatids with a minimum set of purified factors. Nat Cell Biol. 2015;17(8):1014–23. Epub 20150615. doi: 10.1038/ncb3187. PubMed PMID: 26075356.

95. Nonaka N, Kitajima T, Yokobayashi S, Xiao G, Yamamoto M, Grewal SI, et al. Recruitment of cohesin to heterochromatic regions by Swi6/HP1 in fission yeast. Nat Cell Biol. 2002;4(1):89–93. doi: 10.1038/ncb739. PubMed PMID: 11780129.

96. Choppakatla P, Dekker B, Cutts EE, Vannini A, Dekker J, Funabiki H. Linker histone H1.8 inhibits chromatin binding of condensins and DNA topoisomerase II to tune chromosome length and individualization. Elife. 2021;10:37. doi: 10.7554/eLife.68918. PubMed PMID: WOS:000693094000001.

97. Sánchez A, Roguev A, Krogan NJ, Russell P. Genetic Interaction Landscape Reveals Critical Requirements for Schizosaccharomyces pombe Brc1 in DNA Damage Response Mutants. G3 (Bethesda). 2015;5(5):953–62. Epub 2015/03/19. doi: 10.1534/g3.115.017251. PubMed PMID: 25795664; PubMed Central PMCID: PMCPMC4426379.

