## Supplementary Information for "SAGA histone acetyltransferase module facilitates chromatin accessibility to SMC5/6"

#### Materials and methods

#### Yeast two-hybrid analysis

Most of the *S. pombe* and human SMC5/6 Y2H plasmids were prepared previously: pGADT7-spNse1(aa1-232), pGBKT7-spNse1(aa1-232), pACT2-spNse2(aa1-250), pGBKT7-spNse2(aa1-250), pGADT7-spNse3(aa1-328), pGBKT7-spNse3(aa1-328), and pGBKT7-spNse4(aa1-300) in [1]; pACT2-spNse4(aa1-300), pGADT7-hNSE1(aa1-266), pGADT7-hNSE3(aa55-292) and pGADT7-hNSE4b(aa1-333) in [2]; pACT2-spSmc5(aa1-1065), pGBKT7-spSmc5(aa1-1065), pGBKT7-Smc5(aa2-323+732-1065) in [3]; pGADT7-spSmc6(aa1-1140) in [4]; pGADT7-hNSE2(aa1-247), pGADT7-hSMC5(aa4-1101), pGADT7-hSMC6(aa1-1091) in [5]. The TAFA2B cDNA was obtained from the human ORFeome collection (v8.1, http://horfdb.dfci.harvard.edu/hv5/hv8/index.php?page=home) and subcloned into the pACT2 vector in a single step using the Gateway technology. ORF was PCR amplified using primers listed in Suppl. Table ST4 and cloned with In-Fusion cloning system to pGBKT7.

The spNse3(aa1-90) fragment was created by mutating 91$^{st}$ aa to STOP codon (Suppl. Table ST4) in pGADT7-spNse3(aa1-328) [1] using QuickChange Lightning Site-directed mutagenesis kit (Agilent Technologies). The spNse3(aa80-210) and spNse3(aa80-307) fragments were cleaved out by *Nco*I-*Xho*I from respective pTriEx-Nse3 plasmids [2] and inserted into the pGADT7 vector using T4 ligase protocol (New England Biolabs). hNSE4a(aa1-385) was cloned into the pGADT7 vector through *Nde*I-*Xho*I sites by the In-fusion cloning system (Takara). The spSmc5(aa600-835) fragment was amplified by JP295+JP259 primers and inserted into the *Nde*I-*Xho*I sites of pGBKT7 using the T4 ligase protocol.

The other methods are described in the main text.

**References**

1. Sergeant J, Taylor E, Palecek J, Fousteri M, Andrews E, Sweeney S, et al. Composition and architecture of the Schizosaccharomyces pombe Rad18 (Smc5-6) complex. Mol Cell Biol. 2005;25(1):172-84. doi: 10.1128/MCB.25.1.172-184.2005. PubMed PMID: WOS:000226236900015.

2. Hudson JJR, Bednarova K, Kozakova L, Liao CY, Guerineau M, Colnaghi R, et al. Interactions between the Nse3 and Nse4 Components of the SMC5-6 Complex Identify Evolutionarily Conserved Interactions between MAGE and EID Families. Plos One. 2011;6(2):14. doi: 10.1371/journal.pone.0017270. PubMed PMID: WOS:000287764100039.

3. Palecek J, Vidot S, Feng M, Doherty AJ, Lehmann AR. The SMC5-6 DNA repair complex: Bridging of the SMC5-6 heads by the Kleisin, NSE4, and non-Kleisin subunits. J Biol Chem. 2006;281:36952-9. PubMed PMID: 17005570.

4. Zabrady K, Adamus M, Vondrova L, Liao C, Skoupilova H, Novakova M, et al. Chromatin association of the SMC5/6 complex is dependent on binding of its NSE3 subunit to DNA. Nucleic Acids Research. 2016;44(3):1064-79. doi: 10.1093/nar/gkv1021. PubMed PMID: WOS:000371268700017.

5. Adamus M, Lelkes E, Potesil D, Ganji SR, Kolesar P, Zabrady K, et al. Molecular Insights into the Architecture of the Human SMC5/6 Complex. Journal of Molecular Biology. 2020;432(13):3820-37. doi: 10.1016/j.jmb.2020.04.024. PubMed PMID: WOS:000541931500007.

### Figure legends

**Supplementary Figure S1. SMC5/6 genetic interactions**

(**A**) Sensitivity of the original *nse3-R254E* (*nse3/RE*) cells and cells with the cloNAT-*nse3-R254E* construct integrated into the PEM2 background (NAT-*nse3/RE*). Ten-fold serial dilutions of these strains were plated onto YES media containing indicated concentrations of the hydroxyurea (HU) or methyl methane sulfonate (MMS). (**B** and **D**) Tetrad analysis of the diploid fission yeast strains. The original *nse3-R254E* mutant strain was crossed with the indicated mutants and sporulated. Representatives of the tetrad analysis are shown with the double mutants indicated by triangles. SAGA subunits are colour-coded as in panel C. NuA4 subunits are in bold. (**C**) Schematic representation of the SAGA complex with differently coloured subunit modules. Blue, HAT module; orange, DUB module.

**Supplementary Figure S2. Genetic and physical interactions between the SMC5/6 and HAT module**

(**A**) Phenotypes of the SMC5/6 and SAGA mutants. Ten-fold serial dilutions of the indicated strains were plated onto YES media and grown at 25°C (control) or 37°C. The *nse3-R*254E, *smc6*-74, and *smc6*-X mutations enhanced the sensitivity of the Δ*gcn5* mutant to the higher temperature (*ts* phenotype). In contrast, *nse2-SA* did not affect the *ts* phenotype of Δ*gcn5*. Nevertheless, all *smc5/6* mutants enhanced the sensitivity of Δ*gcn5* to the DNA damaging agents (HU or MMS). (**B - D**) The yeast two-hybrid (Y2H) system was used to determine protein-protein interactions between SMC5/6 and SAGA HAT module (Ada2 and Gcn5) subunits. The protein-protein interactions were scored by the growth of yeast PJ69 transformants on the plates without Leu, Trp, and His, containing 3-Amino-1,2,4-triazole. (**B**) The full-length Gal4BD-Gcn5 and Gal4BD-Ada2 (left part) or Gal4AD-Ada2 (right part) hybrid constructs were co-transformed together with the Gal4AD (left panel) or Gal4BD (right panel) full-length SMC5/6 hybrid constructs. The Gal4AD-Nse3 interacted with both Gal4BD-Ada2 and Gal4BD-Gcn5; Gal4BD-Nse2 interacted with Gal4AD-Ada2. (**C**) Detailed Y2H analysis of the above (panel B) interactions and SMC5-Ada2 interaction. Gcn5 interacted with the N-terminal part of Nse3, and Ada2 bound its C-terminal WHB domain. Ada2 bound the N-terminal part of Nse2 and to the arm of SMC5 next to its Nse2-binding interface. Note that the Gal4BD-SMC5 CC arm and Gal4BD-Nse2 full-length constructs self-activated. Therefore, their interactions were scored at higher amino-triazole concentrations. In control experiments, empty pGADT7 and pGBKT7 vectors were used as indicated. (**D**) Human SMC5/6 subunits were tested for their binding to hTADA2B in Y2H. The full-length and hNSE3(aa55-292) Gal4AD hybrid constructs were used. Only Gal4AD-hNSE4a and Gal4AD-hNSE4b construct bound Gal4BD-hTADA2B.

**Supplementary Figure S3. Sensitivity of the SMC5/6 and SAGA strains to genotoxins.**

(**A** and **D**) Ten-fold serial dilutions of the indicated strains were plated onto YES media containing indicated concentrations of HU or MMS. The double mutants were more sensitive than their single mutant counterparts suggesting the non-redundant functions of SMC5/6 and SAGA in DNA repair. (**B**, **C**, and **E**) Comparison of the Nse4-FLAG strains with their parental counterparts. Ten-fold serial dilutions of the Nse4-FLAG strains and their parental counterparts were plated onto YES media with or without indicated concentrations of HU or MMS. The plates were grown at 25°C (or 37°C, if indicated).

**Tables**

**Supplementary Table ST1 - List of genes with significant SGA scores.**

**Supplementary Table ST2 - Summary of significantly enriched GO categories for Biological processes**

**Supplementary Table ST3 - Summary of significantly enriched GO categories for Cellular components**

**Supplementary Table ST4 - List of cloning primers**

**Supplementary Table ST5 - List of yeast strains**

**Supplementary Table STX - List of qPCR primers**

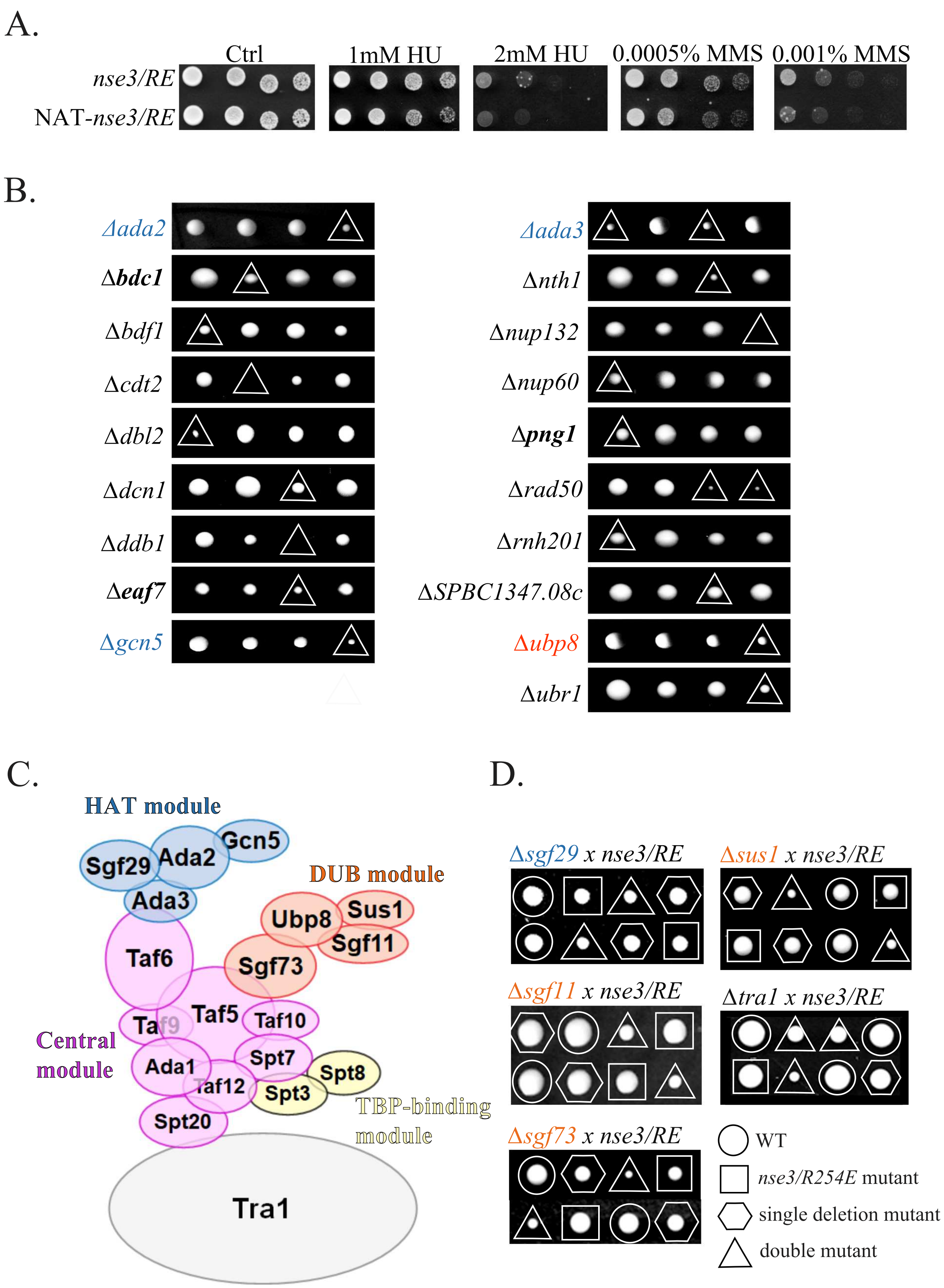
A.
Ctrl
1mM HU
2mM HU
0.0005% MMS
0.001% MMS
nse3/RE
NAT-nse3/RE
B.
Δada2
Δbdc1
Δbdf1
Δcdt2
Δdbl2
Δdcn1
Δddb1
Δeaf7
Δgcn5
Δada3
Δnth1
Δnup132
Δnup60
Δpng1
Δrad50
Δrnh201
ΔSPBC1347.08c
Δubp8
Δubr1
C.
HAT module
Sgf29
Ada2
Gcn5
Ada3
DUB module
Ubp8
Sus1
Sgf11
Sgf73
Taf6
Taf9
Taf5
Taf10
Central module
Ada1
Spt7
Spt8
Taf12
Spt3
TBP-binding module
Spt20
Tra1
D.
Δsgf29 x nse3/RE
Δsus1 x nse3/RE
Δsgf11 x nse3/RE
Δtra1 x nse3/RE
Δsgf73 x nse3/RE
WT
nse3/R254E mutant
single deletion mutant
double mutant

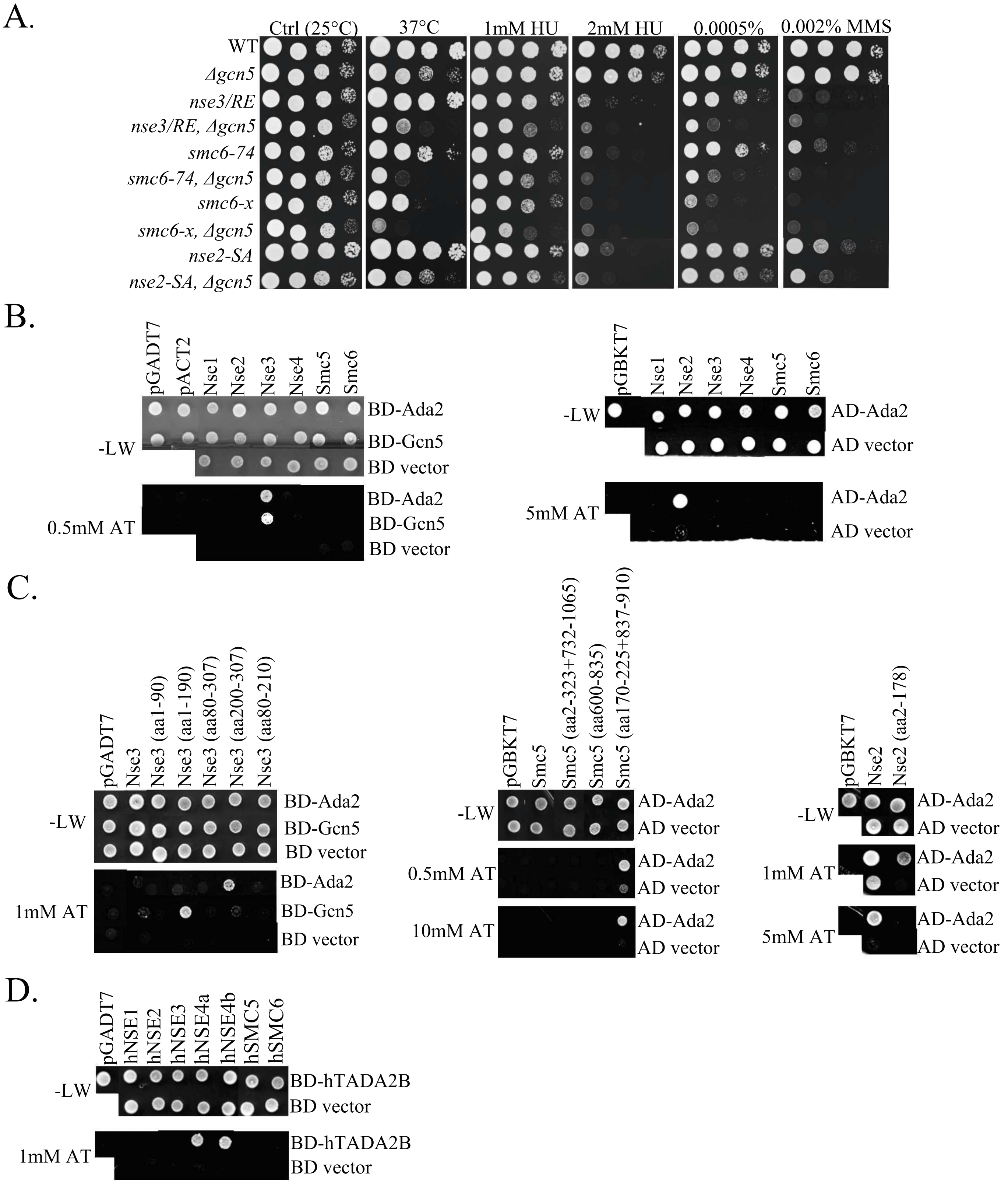
A.
Ctrl (25°C)
37°C
1mM HU
2mM HU
0.0005%
0.002% MMS
WT
Δgcn5
nse3/RE
nse3/RE, Δgcn5
smc6-74
smc6-74, Δgcn5
smc6-x
smc6-x, Δgcn5
nse2-SA
nse2-SA, Δgcn5
B.
pGADT7
pACT2
Nse1
Nse2
Nse3
Nse4
Smc5
Smc6
BD-Ada2
BD-Gcn5
BD vector
-LW
0.5mM AT
pGBKT7
AD-Ada2
AD vector
5mM AT
C.
Nse3 (aa1-90)
Nse3 (aa1-190)
Nse3 (aa80-307)
Nse3 (aa200-307)
Nse3 (aa80-210)
1mM AT
Smc5 (aa2-323+732-1065)
Smc5 (aa600-835)
Smc5 (aa170-225+837-910)
0.5mM AT
10mM AT
Nse2 (aa2-178)
D.
hNSE1
hNSE2
hNSE3
hNSE4a
hNSE4b
hSMC5
hSMC6
BD-hTADA2B

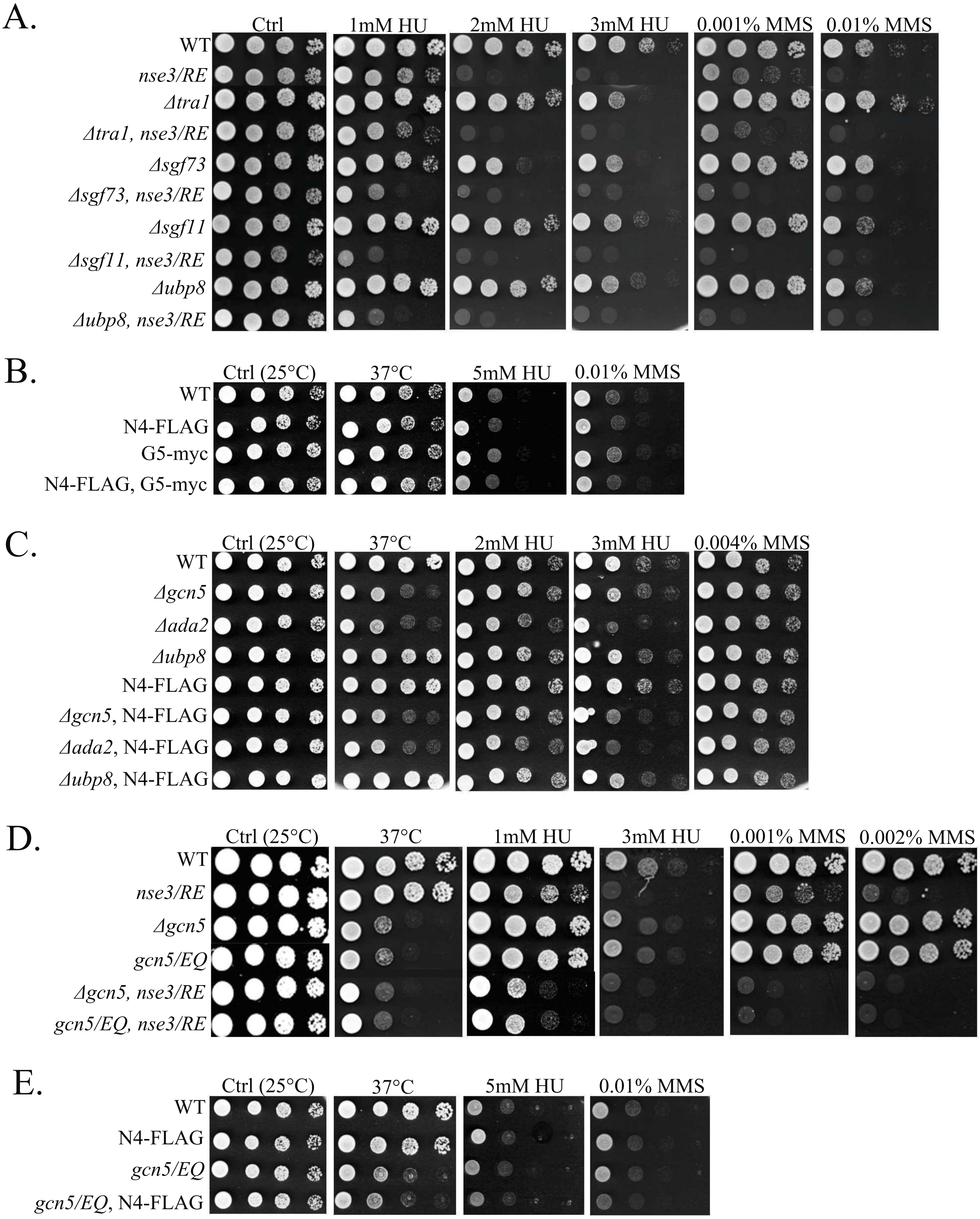
A.
Ctrl
1mM HU
2mM HU
3mM HU
0.001% MMS
0.01% MMS
WT
nse3/RE
Δtra1
Δtra1, nse3/RE
Δsgf73
Δsgf73, nse3/RE
Δsgf11
Δsgf11, nse3/RE
Δubp8
Δubp8, nse3/RE
B.
Ctrl (25°C)
37°C
5mM HU
0.01% MMS
WT
N4-FLAG
G5-myc
N4-FLAG, G5-myc
C.
Ctrl (25°C)
37°C
2mM HU
3mM HU
0.004% MMS
WT
Δgcn5
Δada2
Δubp8
N4-FLAG
Δgcn5, N4-FLAG
Δada2, N4-FLAG
Δubp8, N4-FLAG
D.
Ctrl (25°C)
37°C
1mM HU
3mM HU
0.001% MMS
0.002% MMS
WT
nse3/RE
Δgcn5
gcn5/EQ
Δgcn5, nse3/RE
gcn5/EQ, nse3/RE
E.
Ctrl (25°C)
37°C
5mM HU
0.01% MMS
WT
N4-FLAG
gcn5/EQ
gcn5/EQ, N4-FLAG

**Suppl. Table ST1 - SGA scores**

| POMBASE ID | average score |
|---|---|
| *ada2* | -0,701 |
| ade8 | -0,727 |
| alg14 | -0,429 |
| alp14 | -0,494 |
| **apn2** | -1,071 |
| **ase1** | -0,339 |
| atg22 | -0,422 |
| *bdc1* | -0,381 |
| *bdf1* | -0,393 |
| **brc1** | -0,666 |
| **ccr4** | -0,553 |
| cdr2 | -0,646 |
| ***cdt2*** | -0,441 |
| cgs1 | -1,095 |
| *dbl2* | -0,582 |
| **dbl8** | -0,360 |
| dbs2 | -0,388 |
| **dcd1** | -0,500 |
| *dcn1* | -0,330 |
| ***ddb1*** | -0,462 |
| *eaf7* | -0,337 |
| ect1 | -0,671 |
| **elc1** | -0,303 |
| **fft3** | -0,354 |
| flx1 | -0,792 |
| *gcn5* | -0,545 |
| **hrq1** | -0,760 |
| ies6 | -0,906 |
| lsk1 | -0,486 |
| lsm1 | -0,512 |
| mcp6 | -0,791 |
| mpd2 | -0,695 |
| **mrc1** | -0,571 |
| mtq1 | -0,344 |
| mub1 | -0,436 |
| myh1 | -0,504 |
| nem1 | -0,766 |
| nem2 | -0,527 |
| *ngg1* | -0,446 |
| nrl1 | -0,518 |
| **nse5** | -0,756 |
| **nse6** | -0,684 |
| ***nth1*** | -0,695 |
| ***nup132*** | -0,919 |
| *nup60* | -0,546 |
| *png1* | -0,487 |
| **pnk1** | -0,387 |
| pph3 | -0,393 |
| ppn1 | -0,462 |
| **rad2** | -0,418 |
| **rad26** | -0,535 |
| *rad50* | -0,526 |
| rds1 | -0,497 |
| rei1 | -0,391 |
| **rhp14** | -0,726 |
| **rhp41** | -0,950 |
| *rnh201* | -0,323 |
| ***rnh202*** | -0,393 |
| rsm1 | -0,852 |
| set1 | -0,300 |
| SPAC1039.08 | -0,656 |
| SPAC15E1.02c | -0,516 |
| SPAC513.04 | -1,208 |
| SPAC56F8.02 | -0,515 |
| SPAC8C9.04 | -0,301 |
| SPBC557.02c | -0,650 |
| SPCC23B6.01c | -0,396 |
| SPCC594.01 | -0,762 |
| srb11 | -0,290 |
| **srs2** | -0,842 |
| tdh1 | -0,485 |
| **tdp1** | -0,370 |
| tel1 | -0,285 |
| tsc2 | -0,500 |
| ubi4 | -0,901 |
| *ubp8* | -0,481 |
| *ubr1* | -0,260 |
| ulp2 | -0,712 |
| yox1 | -0,381 |

**bold: negative genetic interactions already described in previous studies**
*italics: genetic interactions verified by tetrad analysis (Suppl. Fig. S1B)*

**Suppl. Table ST2 - GO Biological process**

| p-value | GO-ID | Description | Genes in test set |
|---|---|---|---|
| 3,27E-13 | 6259 | DNA metabolic process | *SPBC3D6.10, SPAC17H9.19C, SPAC26A3.02, SPAC3G6.06C, SPCC24B10.08C, SPAC30D11.07, SPAC11E3.08C, SPAC4H3.05, SPCC553.01C, SPAC1556.01C, SPAC12B10.12C, SPCP31B10.05, SPBC582.06C, SPAC23C11.04C, SPCC23B6.03C, SPBC1347.08C, SPAC17H9.10C, SPBC649.03, SPCC306.04C, SPBC16A3.19, SPAC1952.05, SPBC651.10, SPAC4G9.02, SPAC3G9.08, SPBC582.05C* |
| 1,69E-12 | 6974 | response to DNA damage stimulus | *SPAC9E9.08, SPBC3D6.10, SPAC17H9.19C, SPAC17H9.10C, SPAC26A3.02, SPBC649.03, SPCC306.04C, SPAC3G6.06C, SPBC16A3.19, SPAC694.06C, SPBC651.10, SPAC30D11.07, SPAC11E3.08C, SPAC4H3.05, SPAC1556.01C, SPAC12B10.12C, SPAC3G9.08, SPCP31B10.05, SPAC23C11.04C, SPCC23B6.03C, SPBC582.05C* |
| 9,06E-09 | 6950 | response to stress | *SPAC343.12, SPAC9E9.08, SPBC3D6.10, SPAC17H9.19C, SPAC26A3.02, SPAC2G11.13, SPAC3G6.06C, SPCC24B10.08C, SPAC167.05, SPAC30D11.07, SPAC11E3.08C, SPAC8C9.03, SPAC4H3.05, SPAC1556.01C, SPAC12B10.12C, SPCP31B10.05, SPAC23C11.04C, SPCC23B6.03C, SPAC630.13C, SPCC23B6.01C, SPAC17H9.10C, SPBC649.03, SPCC306.04C, SPAC15E1.02C, SPBC16A3.19, SPAC694.06C, SPBC651.10, SPBC3B8.10C, SPAC3G9.08, SPBC582.05C, SPBC32F12.11* |
| 1,77E-08 | 33554 | cellular response to stress | *SPAC343.12, SPAC9E9.08, SPBC3D6.10, SPAC17H9.19C, SPAC26A3.02, SPAC2G11.13, SPAC3G6.06C, SPAC167.05, SPAC30D11.07, SPAC11E3.08C, SPAC8C9.03, SPAC4H3.05, SPAC1556.01C, SPAC12B10.12C, SPCP31B10.05, SPAC23C11.04C, SPCC23B6.03C, SPAC630.13C, SPCC23B6.01C, SPAC17H9.10C, SPBC649.03, SPCC306.04C, SPAC15E1.02C, SPBC16A3.19, SPAC694.06C, SPBC651.10, SPBC3B8.10C, SPAC3G9.08, SPBC582.05C, SPBC32F12.11* |
| 4,47E-08 | 6281 | DNA repair | *SPBC3D6.10, SPAC17H9.10C, SPAC26A3.02, SPBC649.03, SPAC3G6.06C, SPBC16A3.19, SPAC30D11.07, SPAC4H3.05, SPAC1556.01C, SPAC12B10.12C, SPAC3G9.08, SPCP31B10.05, SPAC23C11.04C, SPBC582.05C* |
| 7,09E-08 | 51716 | cellular response to stimulus | *SPAC343.12, SPAC9E9.08, SPBC3D6.10, SPAC17H9.19C, SPAC26A3.02, SPAC2G11.13, SPAC3G6.06C, SPAC167.05, SPAC30D11.07, SPAC11E3.08C, SPAC8C9.03, SPAC4H3.05, SPAC1556.01C, SPAC12B10.12C, SPCP31B10.05, SPAC23C11.04C, SPCC23B6.03C, SPAC630.13C, SPCC23B6.01C, SPAC17H9.10C, SPBC649.03, SPCC306.04C, SPAC15E1.02C, SPBC16A3.19, SPAC694.06C, SPBC651.10, SPBC3B8.10C, SPAC3G9.08, SPBC582.05C, SPBC32F12.11* |
| 1,14E-07 | 50896 | response to stimulus | *SPAC343.12, SPAC9E9.08, SPBC3D6.10, SPAC17H9.19C, SPAC26A3.02, SPAC2G11.13, SPAC3G6.06C, SPCC24B10.08C, SPAC167.05, SPAC30D11.07, SPAC11E3.08C, SPAC8C9.03, SPAC4H3.05, SPAC1556.01C, SPAC12B10.12C, SPCP31B10.05, SPAC23C11.04C, SPCC23B6.03C, SPAC630.13C, SPCC23B6.01C, SPAC17H9.10C, SPBC649.03, SPCC306.04C, SPAC15E1.02C, SPBC16A3.19, SPAC694.06C, SPBC651.10, SPBC3B8.10C, SPAC3G9.08, SPBC582.05C, SPBC32F12.11* |

| | | | |
|---|---|---|---|
| 1,79E-07 | 75 | cell cycle checkpoint | *SPCC895.07, SPAC1805.04, SPAC9E9.08, SPAC17H9.19C, SPAC2F3.15, SPAPB1A10.09, SPAC1556.01C, SPAC26A3.02, SPCC306.04C, SPCC23B6.03C, SPAC694.06C* |
| 1,66E-06 | 43687 | post-translational protein modification | *SPBC21D10.10, SPAC17H9.19C, SPBC337.08C, SPBC1861.07, SPAC17H9.10C, SPAC13A11.04C, SPBC31F10.10C, SPCC306.04C, SPBC16A3.19, SPAC1952.05, SPBC28F2.10C, SPCC24B10.08C, SPBC3B8.10C, SPAC29B12.05C, SPAC2F3.15, SPAC57A10.02, SPBC26H8.05C, SPBC19C7.02, SPBC839.03C, SPAC17A5.07C, SPCC23B6.03C* |
| 2,04E-06 | 77 | DNA damage checkpoint | *SPAC9E9.08, SPAC17H9.19C, SPAC1556.01C, SPAC26A3.02, SPCC306.04C, SPCC23B6.03C, SPAC694.06C* |
| 2,85E-06 | 42770 | DNA damage response, signal transduction | *SPAC9E9.08, SPAC17H9.19C, SPAC1556.01C, SPAC26A3.02, SPCC306.04C, SPCC23B6.03C, SPAC694.06C* |
| 6,16E-06 | 90304 | nucleic acid metabolic process | *SPCC74.02C, SPBC3D6.10, SPAC17H9.19C, SPAC26A3.02, SPBC29A10.10C, SPAC3G6.06C, SPCC24B10.08C, SPAC30D11.07, SPAC11E3.08C, SPAC4H3.05, SPCC553.01C, SPAC1556.01C, SPAC12B10.12C, SPCP31B10.05, SPBC582.06C, SPAC23C11.04C, SPCC23B6.03C, SPBC12D12.06, SPBC1861.07, SPBC1347.08C, SPAC17H9.10C, SPAC13A11.04C, SPBC649.03, SPCC306.04C, SPBC16A3.19, SPAC1952.05, SPBC28F2.10C, SPBC651.10, SPBC3D6.08C, SPAC4G9.02, SPCC31H12.08C, SPAC3G9.08, SPBC582.05C* |
| 7,32E-06 | 51276 | chromosome organization | *SPBC21D10.10, SPAC13A11.04C, SPCC1450.02, SPBC649.03, SPCC306.04C, SPAC222.04C, SPAC3G6.06C, SPBC16A3.19, SPAC1952.05, SPBC28F2.10C, SPCC24B10.08C, SPAC25A8.01C, SPCC553.01C, SPAC1556.01C, SPAC17A5.07C, SPBC582.06C, SPCC23B6.03C, SPBC582.05C* |
| 9,34E-06 | 51052 | regulation of DNA metabolic process | *SPBC651.10, SPAC9E9.08, SPAC17H9.19C, SPAC11E3.08C, SPAC17H9.10C, SPAC3G9.08, SPBC582.05C, SPAC694.06C* |
| 1,21E-05 | 6139 | nucleobase, nucleoside, nucleotide and nucleic acid metabolic process | *SPCC74.02C, SPBC3D6.10, SPAC17H9.19C, SPAC26A3.02, SPBC29A10.10C, SPAC3G6.06C, SPCC24B10.08C, SPAC30D11.07, SPAC11E3.08C, SPAC4H3.05, SPCC553.01C, SPAC1556.01C, SPAC12B10.12C, SPCP31B10.05, SPBC582.06C, SPAC23C11.04C, SPCC23B6.03C, SPBC12D12.06, SPBC14F5.09C, SPBC1861.07, SPBC1347.08C, SPAC17H9.10C, SPAC13A11.04C, SPBC649.03, SPCC306.04C, SPBC16A3.19, SPAC1952.05, SPBC28F2.10C, SPBC2G2.13C, SPBC651.10, SPBC3D6.08C, SPAC4G9.02, SPCC31H12.08C, SPAC3G9.08, SPBC582.05C* |
| 1,98E-05 | 31570 | DNA integrity checkpoint | *SPAC9E9.08, SPAC17H9.19C, SPAC1556.01C, SPAC26A3.02, SPCC306.04C, SPCC23B6.03C, SPAC694.06C* |
| 2,50E-05 | 16573 | histone acetylation | *SPCC24B10.08C, SPBC21D10.10, SPAC13A11.04C, SPBC16A3.19, SPAC1952.05, SPBC28F2.10C* |
| 3,35E-05 | 6464 | protein modification process | *SPBC21D10.10, SPAC17H9.19C, SPBC337.08C, SPBC1861.07, SPAC17H9.10C, SPAC13A11.04C, SPBC31F10.10C, SPCC306.04C, SPBC16A3.19, SPAC1952.05, SPBC28F2.10C, SPCC24B10.08C, SPBC3B8.10C, SPAC29B12.05C, SPAC2F3.15, SPAC57A10.02, SPBC26H8.05C, SPBC19C7.02, SPBC839.03C, SPAC17A5.07C, SPCC23B6.03C, SPAC5D6.06C* |

| | | | |
|---|---|---|---|
| 6,27E-05 | 51327 | M phase of meiotic cell cycle | *SPCC24B10.08C, SPAC167.05, SPBC902.03, SPBC337.08C, SPAC8C9.04, SPAPB1A10.09, SPCC553.01C, SPAC1556.01C, SPBC31F10.10C, SPBC582.06C, SPAC1952.05* |
| 6,27E-05 | 7126 | meiosis | *SPCC24B10.08C, SPAC167.05, SPBC902.03, SPBC337.08C, SPAC8C9.04, SPAPB1A10.09, SPCC553.01C, SPAC1556.01C, SPBC31F10.10C, SPBC582.06C, SPAC1952.05* |
| 6,62E-05 | 51726 | regulation of cell cycle | *SPAC1805.04, SPAC9E9.08, SPAC17H9.19C, SPAC17H9.10C, SPAC26A3.02, SPCC306.04C, SPAC694.06C, SPCC895.07, SPAC2F3.15, SPAC8C9.03, SPAC57A10.02, SPAPB1A10.09, SPAC1556.01C, SPCC23B6.03C* |
| 7,36E-05 | 34641 | cellular nitrogen compound metabolic process | *SPCC74.02C, SPBC3D6.10, SPAC17H9.19C, SPAC26A3.02, SPBC29A10.10C, SPAC3G6.06C, SPCC24B10.08C, SPAC30D11.07, SPAC11E3.08C, SPAC4H3.05, SPCC553.01C, SPAC1556.01C, SPAC12B10.12C, SPCP31B10.05, SPBC582.06C, SPAC23C11.04C, SPCC23B6.03C, SPBC12D12.06, SPBC14F5.09C, SPBC1861.07, SPBC1347.08C, SPAC17H9.10C, SPAC13A11.04C, SPBC649.03, SPCC306.04C, SPBC16A3.19, SPAC1952.05, SPBC28F2.10C, SPBC2G2.13C, SPBC651.10, SPAC15E1.05C, SPBC3D6.08C, SPAC4G9.02, SPCC31H12.08C, SPAC1039.08, SPAC3G9.08, SPBC582.05C* |
| 7,68E-05 | 51321 | meiotic cell cycle | *SPCC24B10.08C, SPAC167.05, SPBC902.03, SPBC337.08C, SPAC8C9.04, SPAPB1A10.09, SPCC553.01C, SPAC1556.01C, SPBC31F10.10C, SPBC582.06C, SPAC1952.05* |
| 8,54E-05 | 6807 | nitrogen compound metabolic process | *SPCC74.02C, SPBC3D6.10, SPAC17H9.19C, SPAC26A3.02, SPBC29A10.10C, SPAC3G6.06C, SPCC24B10.08C, SPAC30D11.07, SPAC11E3.08C, SPAC4H3.05, SPCC553.01C, SPAC1556.01C, SPAC12B10.12C, SPCP31B10.05, SPBC582.06C, SPAC23C11.04C, SPCC23B6.03C, SPBC12D12.06, SPBC14F5.09C, SPBC1861.07, SPBC1347.08C, SPAC17H9.10C, SPAC13A11.04C, SPBC649.03, SPCC306.04C, SPBC16A3.19, SPAC1952.05, SPBC28F2.10C, SPBC2G2.13C, SPBC651.10, SPAC15E1.05C, SPBC3D6.08C, SPAC4G9.02, SPCC31H12.08C, SPAC1039.08, SPAC3G9.08, SPBC582.05C* |
| 1,15E-04 | 6260 | DNA replication | *SPBC651.10, SPAC4G9.02, SPAC17H9.19C, SPAC11E3.08C, SPBC1347.08C, SPAC17H9.10C, SPAC4H3.05, SPAC1556.01C, SPAC3G6.06C* |
| 1,20E-04 | 6338 | chromatin remodeling | *SPCC24B10.08C, SPBC21D10.10, SPAC25A8.01C, SPAC13A11.04C, SPCC1450.02, SPCC306.04C, SPAC222.04C, SPBC16A3.19, SPAC1952.05, SPBC28F2.10C* |
| 1,41E-04 | 16569 | covalent chromatin modification | *SPCC24B10.08C, SPBC21D10.10, SPAC13A11.04C, SPCC306.04C, SPCC23B6.03C, SPBC16A3.19, SPAC1952.05, SPBC28F2.10C* |
| 1,41E-04 | 16570 | histone modification | *SPCC24B10.08C, SPBC21D10.10, SPAC13A11.04C, SPCC306.04C, SPCC23B6.03C, SPBC16A3.19, SPAC1952.05, SPBC28F2.10C* |
| 1,42E-04 | 6473 | protein amino acid acetylation | *SPCC24B10.08C, SPBC21D10.10, SPAC13A11.04C, SPBC16A3.19, SPAC1952.05, SPBC28F2.10C* |
| 1,50E-04 | 16568 | chromatin modification | *SPCC24B10.08C, SPBC21D10.10, SPAC25A8.01C, SPAC13A11.04C, SPCC1450.02, SPCC306.04C, SPAC222.04C, SPCC23B6.03C, SPBC16A3.19, SPAC1952.05, SPBC28F2.10C* |

| | | | |
|---|---|---|---|
| 1,95E-04 | 50789 | regulation of biological process | *SPBC21B10.13C, SPAC9E9.08, SPAC17H9.19C, SPAC26A3.02, SPCC24B10.08C, SPAC11E3.08C, SPAC25A8.01C, SPAC8C9.03, SPAC57A10.02, SPAC1556.01C, SPBC19C7.02, SPCC23B6.03C, SPAC630.13C, SPAC1805.04, SPBC1861.07, SPAC17H9.10C, SPCC306.04C, SPBC16A3.19, SPAC694.06C, SPAC1952.05, SPCC895.07, SPBC651.10, SPBC3B8.10C, SPBC902.03, SPAC2F3.15, SPAPB1A10.09, SPBC26H8.05C, SPAC3G9.08, SPBC839.03C, SPBC582.05C, SPBC32F12.11* |
| 2,17E-04 | 43412 | macromolecule modification | *SPBC21D10.10, SPAC17H9.19C, SPBC337.08C, SPBC1861.07, SPAC17H9.10C, SPAC13A11.04C, SPBC31F10.10C, SPCC306.04C, SPBC16A3.19, SPAC1952.05, SPBC28F2.10C, SPCC24B10.08C, SPBC3B8.10C, SPAC29B12.05C, SPAC2F3.15, SPAC57A10.02, SPBC26H8.05C, SPBC19C7.02, SPBC839.03C, SPAC17A5.07C, SPCC23B6.03C, SPAC5D6.06C* |
| 2,61E-04 | 6284 | base-excision repair | *SPAC30D11.07, SPBC3D6.10, SPAC26A3.02* |
| 3,08E-04 | 43543 | protein amino acid acylation | *SPCC24B10.08C, SPBC21D10.10, SPAC13A11.04C, SPBC16A3.19, SPAC1952.05, SPBC28F2.10C* |
| 3,65E-04 | 6325 | chromatin organization | *SPCC24B10.08C, SPBC21D10.10, SPAC25A8.01C, SPAC13A11.04C, SPCC1450.02, SPCC306.04C, SPAC222.04C, SPCC23B6.03C, SPBC16A3.19, SPAC1952.05, SPBC28F2.10C* |
| 3,69E-04 | 80135 | regulation of cellular response to stress | *SPAC17H9.10C, SPAC3G9.08, SPBC32F12.11* |
| 3,69E-04 | 71478 | cellular response to radiation | *SPAC17H9.10C, SPAC4H3.05, SPAC26A3.02* |
| 3,69E-04 | 71482 | cellular response to light stimulus | *SPAC17H9.10C, SPAC4H3.05, SPAC26A3.02* |
| 3,69E-04 | 34644 | cellular response to UV | *SPAC17H9.10C, SPAC4H3.05, SPAC26A3.02* |
| 4,62E-04 | 23046 | signaling process | *SPAC1805.04, SPAC9E9.08, SPAC17H9.19C, SPAC26A3.02, SPCC306.04C, SPAC694.06C, SPCC895.07, SPAC8C9.03, SPAC57A10.02, SPBC26H8.05C, SPAC1556.01C, SPCC23B6.03C, SPBC32F12.11, SPAC630.13C* |
| 4,62E-04 | 23060 | signal transmission | *SPAC1805.04, SPAC9E9.08, SPAC17H9.19C, SPAC26A3.02, SPCC306.04C, SPAC694.06C, SPCC895.07, SPAC8C9.03, SPAC57A10.02, SPBC26H8.05C, SPAC1556.01C, SPCC23B6.03C, SPBC32F12.11, SPAC630.13C* |
| 4,62E-04 | 7165 | signal transduction | *SPAC1805.04, SPAC9E9.08, SPAC17H9.19C, SPAC26A3.02, SPCC306.04C, SPAC694.06C, SPCC895.07, SPAC8C9.03, SPAC57A10.02, SPBC26H8.05C, SPAC1556.01C, SPCC23B6.03C, SPBC32F12.11,* |
| 5,02E-04 | 80134 | regulation of response to stress | *SPAC17H9.10C, SPAC3G9.08, SPBC32F12.11* |
| 5,85E-04 | 8156 | negative regulation of DNA replication | *SPBC651.10, SPAC9E9.08, SPAC17H9.19C, SPAC11E3.08C, SPAC694.06C* |
| 6,16E-04 | 70647 | protein modification by small protein conjugation or removal | *SPAC17H9.19C, SPBC337.08C, SPBC1861.07, SPAC17H9.10C, SPAC13A11.04C, SPBC19C7.02, SPBC31F10.10C, SPBC839.03C, SPAC17A5.07C* |
| 6,62E-04 | 43137 | DNA replication, removal of RNA primer | *SPAC4G9.02, SPBC1347.08C, SPAC3G6.06C* |
| 7,50E-04 | 6289 | nucleotide-excision repair | *SPAC17H9.10C, SPAC12B10.12C, SPBC649.03, SPAC23C11.04C* |

| | | | |
|---|---|---|---|
| 7,59E-04 | 44265 | cellular macromolecule catabolic process | *SPAC30D11.07, SPBC3D6.08C, SPAC4G9.02, SPAC17H9.19C, SPBC1861.07, SPBC1347.08C, SPAC17H9.10C, SPAC13A11.04C, SPCC31H12.08C, SPBC19C7.02, SPAC23C11.04C, SPAC3G6.06C* |
| 7,84E-04 | 279 | M phase | *SPCC24B10.08C, SPAC167.05, SPCC895.07, SPBC902.03, SPBC337.08C, SPAC8C9.04, SPAPB1A10.09, SPCC553.01C, SPAC1556.01C, SPBC31F10.10C, SPBC582.06C, SPAC1952.05* |
| 7,95E-04 | 65007 | biological regulation | *SPBC21B10.13C, SPAC9E9.08, SPAC17H9.19C, SPAC26A3.02, SPAC3G6.06C, SPCC24B10.08C, SPAC11E3.08C, SPAC25A8.01C, SPAC8C9.03, SPAC57A10.02, SPCC553.01C, SPAC1556.01C, SPBC19C7.02, SPCC23B6.03C, SPAC630.13C, SPAC1805.04, SPBC1861.07, SPAC17H9.10C, SPCC306.04C, SPBC16A3.19, SPAC694.06C, SPAC1952.05, SPCC895.07, SPBC651.10, SPBC3B8.10C, SPBC902.03, SPAC2F3.15, SPAPB1A10.09, SPBC26H8.05C, SPAC3G9.08, SPBC839.03C, SPBC582.05C, SPBC32F12.11* |
| 7,99E-04 | 23052 | signaling | *SPAC1805.04, SPAC9E9.08, SPAC17H9.19C, SPAC26A3.02, SPCC306.04C, SPAC694.06C, SPCC895.07, SPAC8C9.03, SPAC57A10.02, SPBC26H8.05C, SPAC1556.01C, SPCC23B6.03C, SPBC32F12.11, SPAC630.13C* |
| 8,51E-04 | 9411 | response to UV | *SPAC17H9.10C, SPAC4H3.05, SPAC26A3.02* |
| 8,51E-04 | 9416 | response to light stimulus | *SPAC17H9.10C, SPAC4H3.05, SPAC26A3.02* |
| 8,51E-04 | 48583 | regulation of response to stimulus | *SPAC17H9.10C, SPAC3G9.08, SPBC32F12.11* |
| 8,83E-04 | 22403 | cell cycle phase | *SPAC9E9.08, SPBC337.08C, SPBC31F10.10C, SPAC1952.05, SPCC24B10.08C, SPAC167.05, SPCC895.07, SPBC902.03, SPAC8C9.04, SPAPB1A10.09, SPCC553.01C, SPAC1556.01C, SPBC582.06C* |
| 9,44E-04 | 51053 | negative regulation of DNA metabolic process | *SPBC651.10, SPAC9E9.08, SPAC17H9.19C, SPAC11E3.08C, SPAC694.06C* |
| 1,07E-03 | 9314 | response to radiation | *SPAC17H9.10C, SPAC4H3.05, SPAC26A3.02* |
| 1,07E-03 | 71214 | cellular response to abiotic stimulus | *SPAC17H9.10C, SPAC4H3.05, SPAC26A3.02* |
| 1,07E-03 | 33567 | DNA replication, Okazaki fragment | *SPAC4G9.02, SPBC1347.08C, SPAC3G6.06C* |
| 1,19E-03 | 6261 | DNA-dependent DNA replication | *SPBC651.10, SPAC4G9.02, SPAC17H9.19C, SPAC11E3.08C, SPBC1347.08C, SPAC4H3.05, SPAC3G6.06C* |
| 1,32E-03 | 6273 | lagging strand elongation | *SPAC4G9.02, SPBC1347.08C, SPAC3G6.06C* |
| 1,32E-03 | 31573 | intra-S DNA damage checkpoint | *SPAC9E9.08, SPAC1556.01C, SPAC694.06C* |
| 1,33E-03 | 6275 | regulation of DNA replication | *SPBC651.10, SPAC9E9.08, SPAC17H9.19C, SPAC11E3.08C, SPAC694.06C* |
| 1,33E-03 | 32200 | telomere organization | *SPCC553.01C, SPAC1556.01C, SPCC306.04C, SPCC23B6.03C, SPAC3G6.06C* |
| 1,33E-03 | 723 | telomere maintenance | *SPCC553.01C, SPAC1556.01C, SPCC306.04C, SPCC23B6.03C, SPAC3G6.06C* |
| 1,33E-03 | 60249 | anatomical structure homeostasis | *SPCC553.01C, SPAC1556.01C, SPCC306.04C, SPCC23B6.03C, SPAC3G6.06C* |
| 1,49E-03 | 9057 | macromolecule catabolic process | *SPAC30D11.07, SPBC3D6.08C, SPAC4G9.02, SPAC17H9.19C, SPBC1861.07, SPBC1347.08C, SPAC17H9.10C, SPAC13A11.04C, SPCC31H12.08C, SPBC19C7.02, SPAC23C11.04C, SPAC3G6.06C* |

| | | | |
|---|---|---|---|
| 1,51E-03 | 22402 | cell cycle process | *SPAC9E9.08, SPBC337.08C, SPBC31F10.10C, SPAC1952.05, SPCC24B10.08C, SPAC167.05, SPCC895.07,* |
| 1,74E-03 | 50794 | regulation of cellular process | *SPBC21B10.13C, SPAC9E9.08, SPAC17H9.19C, SPAC26A3.02, SPCC24B10.08C, SPAC11E3.08C, SPAC25A8.01C, SPAC8C9.03, SPAC57A10.02, SPAC1556.01C, SPCC23B6.03C, SPAC630.13C, SPAC1805.04, SPBC1861.07, SPAC17H9.10C, SPCC306.04C, SPBC16A3.19, SPAC694.06C, SPAC1952.05, SPCC895.07, SPBC651.10, SPAC2F3.15, SPAPB1A10.09, SPBC26H8.05C, SPAC3G9.08, SPBC839.03C, SPBC582.05C, SPBC32F12.11* |
| 2,19E-03 | 6282 | regulation of DNA repair | *SPAC17H9.10C, SPAC3G9.08* |
| 2,19E-03 | 31565 | cytokinesis checkpoint | *SPAC2F3.15, SPAPB1A10.09* |
| 2,49E-03 | 23034 | intracellular signaling pathway | *SPAC9E9.08, SPAC17H9.19C, SPAC1556.01C, SPAC26A3.02, SPCC306.04C, SPCC23B6.03C, SPBC32F12.11, SPAC630.13C, SPAC694.06C* |
| 2,76E-03 | 80090 | regulation of primary metabolic process | *SPBC21B10.13C, SPAC9E9.08, SPAC17H9.19C, SPBC1861.07, SPAC17H9.10C, SPBC16A3.19, SPAC694.06C, SPAC1952.05, SPCC24B10.08C, SPBC651.10, SPBC3B8.10C, SPBC902.03, SPAC11E3.08C, SPAC2F3.15, SPAC8C9.03, SPAC3G9.08, SPBC839.03C, SPBC582.05C, SPAC630.13C* |
| 2,98E-03 | 6996 | organelle organization | *SPBC21D10.10, SPAC1805.04, SPAC13A11.04C, SPCC1450.02, SPBC649.03, SPCC306.04C, SPAC222.04C, SPAC3G6.06C, SPBC16A3.19, SPAC1952.05, SPBC28F2.10C, SPCC24B10.08C, SPCC895.07, SPBC3B8.10C, SPBC902.03, SPAC29B12.05C, SPAC25A8.01C, SPAPB1A10.09, SPCC553.01C, SPAC1556.01C, SPAC17A5.07C, SPBC582.06C, SPCC23B6.03C, SPBC582.05C* |
| 3,63E-03 | 44093 | positive regulation of molecular function | *SPAC17H9.19C, SPBC839.03C, SPAC630.13C* |
| 3,63E-03 | 43085 | positive regulation of catalytic activity | *SPAC17H9.19C, SPBC839.03C, SPAC630.13C* |
| 3,72E-03 | 23033 | signaling pathway | *SPAC9E9.08, SPAC17H9.19C, SPAC1556.01C, SPAC26A3.02, SPCC306.04C, SPCC23B6.03C, SPBC32F12.11, SPAC630.13C, SPAC694.06C* |
| 4,03E-03 | 32446 | protein modification by small protein conjugation | *SPAC17H9.19C, SPBC337.08C, SPBC1861.07, SPAC17H9.10C, SPBC19C7.02, SPBC31F10.10C, SPBC839.03C* |
| 4,50E-03 | 51443 | positive regulation of ubiquitin-protein ligase activity | *SPAC17H9.19C, SPBC839.03C* |
| 4,50E-03 | 31398 | positive regulation of protein ubiquitination | *SPAC17H9.19C, SPBC839.03C* |
| 4,61E-03 | 7049 | cell cycle | *SPAC9E9.08, SPBC337.08C, SPBC31F10.10C, SPAC1952.05, SPCC24B10.08C, SPAC167.05, SPCC895.07, SPBC902.03, SPAC2F3.15, SPAC8C9.04, SPAPB1A10.09, SPCC553.01C, SPAC1556.01C, SPBC582.06C* |
| 5,33E-03 | 19222 | regulation of metabolic process | *SPBC21B10.13C, SPAC9E9.08, SPAC17H9.19C, SPBC1861.07, SPAC17H9.10C, SPBC16A3.19, SPAC694.06C, SPAC1952.05, SPCC24B10.08C, SPBC651.10, SPBC3B8.10C, SPBC902.03, SPAC11E3.08C,* |

| | | | |
|---|---|---|---|
| 5,67E-03 | 7131 | reciprocal meiotic recombination | *SPCC24B10.08C, SPAC1556.01C, SPBC582.06C, SPAC1952.05* |
| 5,95E-03 | 51351 | positive regulation of ligase activity | *SPAC17H9.19C, SPBC839.03C* |
| 5,95E-03 | 46579 | positive regulation of Ras protein signal transduction | *SPAC2F3.15, SPAC25A8.01C* |
| 5,95E-03 | 46890 | regulation of lipid biosynthetic process | *SPBC3B8.10C, SPBC902.03* |
| 5,95E-03 | 51057 | positive regulation of small GTPase mediated signal transduction | *SPAC2F3.15, SPAC25A8.01C* |
| 6,01E-03 | 70646 | protein modification by small protein removal | *SPBC337.08C, SPAC13A11.04C, SPAC17A5.07C* |
| 6,26E-03 | 19219 | regulation of nucleobase, nucleoside, nucleotide and nucleic acid metabolic process | *SPBC21B10.13C, SPAC9E9.08, SPAC17H9.19C, SPBC1861.07, SPAC17H9.10C, SPBC16A3.19, SPAC694.06C, SPAC1952.05, SPCC24B10.08C, SPBC651.10, SPAC11E3.08C, SPAC2F3.15, SPAC3G9.08, SPBC582.05C, SPAC630.13C* |
| 6,53E-03 | 6401 | RNA catabolic process | *SPBC3D6.08C, SPAC4G9.02, SPBC1347.08C, SPCC31H12.08C, SPAC3G6.06C* |
| 6,87E-03 | 51171 | regulation of nitrogen compound metabolic process | *SPBC21B10.13C, SPAC9E9.08, SPAC17H9.19C, SPBC1861.07, SPAC17H9.10C, SPBC16A3.19, SPAC694.06C, SPAC1952.05, SPCC24B10.08C, SPBC651.10, SPAC11E3.08C, SPAC2F3.15, SPAC3G9.08, SPBC582.05C, SPAC630.13C* |

**Suppl. Table ST3 - GO Cellular Component**

| p-value | GO-ID | Description | Genes in test set |
|---|---|---|---|
| 9,86E-07 | 123 | histone acetyltransferase complex | *ada2, ada3, bdc1, eaf7, gcn5, png1, ubp8* |
| 3,38E-06 | 5634 | nucleus | *ada2, ada3, ade8, alp14, apn2, bdc1, bdf1, brc1, ccr4, cdr2, cdt2, cgs1, dbl2, dbl8, dcd1, dcn1, ddb1, eaf7, ect1, fft3, gcn5, hrq1, ies6, lsk1, lsm1, mcp6, mrc1, mub1, myh1, nrl1, nse5, nse6, nth1, nup132, nup60, png1, pnk1, ppe2, ppn1, rad2, rad26, rad50, rhp14, rhp41, rnh201, rsm1, set1, SPAC1039.08, SPBC1347.08C, SPBC1861.07, SPBC557.02C, SPCC23B6.01C, srb11, srs2, tdp1, tsc2, ubp8, ubr1, ulp2, yox1* |
| 6,71E-05 | 43234 | protein complex | *ada2, ada3, alp14, ase1, bdc1, bdf1, ccr4, cdr2, cgs1, ddb1, eaf7, fft3, gcn5, ies6, lsk1, lsm1, mrc1, nem1, nse5, nse6, nup132, nup60, png1, rad50, rhp14, rhp41, rnh201, set1, SPBC1347.08C, SPBC1861.07, SPBC902.03, srb11, tsc2, ubp8, yox1* |
| 1,13E-04 | 44451 | nucleoplasm part | *ada2, ada3, bdc1, ccr4, eaf7, gcn5, lsk1, png1, set1, SPBC1861.07, srb11, ubp8, yox1* |
| 1,58E-04 | 70461 | SAGA-type complex | *ada2, ada3, gcn5, ubp8* |
| 1,58E-04 | 124 | SAGA complex | *ada2, ada3, gcn5, ubp8* |
| 1,84E-04 | 5654 | nucleoplasm | *ada2, ada3, bdc1, ccr4, eaf7, gcn5, lsk1, png1, set1, SPBC1861.07, srb11, ubp8, yox1* |
| 3,78E-04 | 44427 | chromosomal part | *ada2, ada3, alp14, bdc1, bdf1, eaf7, fft3, gcn5, ies6, mrc1, nse5, nse6, rad2, rad50, set1, tel1* |
| 5,61E-04 | 790 | nuclear chromatin | *ada2, ada3, bdc1, bdf1, eaf7, fft3, gcn5, ies6, mrc1, rad50, set1* |
| 6,07E-04 | 5694 | chromosome | *ada2, ada3, alp14, bdc1, bdf1, eaf7, fft3, gcn5, ies6, mrc1, nse5, nse6, rad2, rad50, set1, tel1* |
| 6,70E-04 | 32299 | ribonuclease H2 complex | *rnh201, SPBC1347.08C* |
| 8,53E-04 | 35267 | NuA4 histone acetyltransferase complex | *bdc1, eaf7, png1* |
| 1,07E-03 | 43189 | H4/H2A histone acetyltransferase complex | *bdc1, eaf7, png1* |
| 1,32E-03 | 44454 | nuclear chromosome part | *ada2, ada3, alp14, bdc1, bdf1, eaf7, fft3, gcn5, ies6, mrc1, rad2, rad50, set1* |
| 1,76E-03 | 785 | chromatin | *ada2, ada3, bdc1, bdf1, eaf7, fft3, gcn5, ies6, mrc1, rad50, set1* |
| 2,12E-03 | 228 | nuclear chromosome | *ada2, ada3, alp14, bdc1, bdf1, eaf7, fft3, gcn5, ies6, mrc1, rad2, rad50, set1* |
| 3,15E-03 | 109 | nucleotide-excision repair complex | *ddb1, rhp14, rhp41* |
| 5,25E-03 | 5819 | spindle | *alp14, ase1, cdr2, dbl2, mcp6, mub1, nse5, rhp41, SPAC56F8.02, srs2* |
| 5,96E-03 | 30915 | Smc5/6 complex | *nse5, nse6* |

**Suppl. Table ST4 - Cloning primers**

| | | | |
|---|---|---|---|
| pGADT7-spAda2 | LJ121 | fw | 5' GGAGGCCAGTGAATTCATGCCTCCTCAAAAGTATCATTGC 3' |
| | LJ122 | rev | 5' CACCCGGGTGGAATTCTTAGGTTGGAGCGCCAATC 3' |
| pGBKT7-spAda2 | LJ119 | fw | 5' CATGGAGGCCGAATTCATGCCTCCTCAAAAGTATCATTGC 3' |
| | LJ120 | rev | 5' GGATCCCCGGGAATTCTTAGGTTGGAGCGCCAATC 3' |
| pGBKT7-spGcn5 | LJ101 | fw | 5' GAATTCCCGGGGATCCCATCGAATTCGTTAAATGATCAAAC 3' |
| | LJ102 | rev | 5' GCAGGTCGACGGATCCTTAATCGGCTAAGTGTGAATA 3' |
| pGBKT7-hTADA2B | LJ209 | fw | 5' CATGGAGGCCGAATTCATGGCTGCCCACGTTG 3' |
| | LJ210 | rev | 5' GGATCCCCGGGAATTCTCAAGACGCGTCCCTGG 3' |
| pGADT7-spNse3 (aa 1-90) | oPK171 | fw | 5' GAACGGAATGAAACAGACGCAATAAATTTTTAACTATTGGTACGAAA 3' |
| | oPK172 | rev | 5' TTTCGTACCAATAGTTAAAAATTTATTGCGTCTGTTTCATTCCGTTC 3' |
| pGADT7-spNse3 (aa 1-190) | oPK173 | fw | 5' GTGTTGAGGTCTACCCTTCCTTAGGAACTTCAAAAGGATTC 3' |
| | oPK174 | rev | 5' GAATCCTTTTGAAGTTCCTAAGGAAGGGTAGACCTCAACAC 3' |
| pGADT7-hNSE4a | oLV111 | fw | 5' CAGATTACGCTCATATGTCTGGGGACAGCAGC 3' |
| | oLV112 | rev | 5' TCATCTGCAGCTCGAGTCAAGCACTTGGCTTCTGC 3' |
| pGBKT7-spSMC5 (aa170-225+837-910) | JP125 | fw | 5' GGGACCATGGCACATGAGAAGCTTATTGATTTA 3' |
| | JP126 | rev | 5' GGAGCGGCCGCTAGTGATGGTGATGGTGATGAGAAAATCTATCGCTAATACATTG 3' |
| pGBKT7-spSMC5 (aa600-835) | JP295 | fw | 5' GACGACGACAAGATGTACGGTGATAGAGAAAT 3' |
| | JP259 | rev | 5' GAGGAGAAGCCCGGTTCACACTTCCGAACTAGTAGCAAA 3' |
| pGBKT7-spSMC6 (aa1-1140) | oLV465 | fw | 5' AGGAGGACCTGCATATGATGACTACAGAGCTTACTAATGTATCT 3' |
| | oLV466 | rev | 5' GCCGCTGCAGGTCGACTCATGGCGCCGTAGA 3' |
| pAW8-spNse3, *Sph* I to *Xho* I mutation | LJ48 | fw | 5' CATACATTATACGAAGTTATCTCGAGGTGAATACGGTAGATAC 3' |
| | LJ49 | rev | 5' GTATCTACCGTATTCACCTCGAGATAACTTCGTATAATGTATG 3' |
| pAW8-cloNAT-spNse3, cloNAT casette insertion | LJ42 | fw | 5' ACGAAGTTATCTCGAGTGCTTTCTCAGGTATAGTATGAGGTC 3' |
| | LJ43 | rev | 5' CCGTATTCACCTCGATGGAATTCGAGCTCGTTTAAACTGGATG 3' |
| pAW8-cloNAT-spNse3, 5'-end extension | LJ44 | fw | 5' ACGAAGTTATCTCGAGACTAGTTAATTGACAGCTCAGCAATACAAC 3' |
| | LJ45 | rev | 5' CCTGAGAAAGCACTCGAGTAACATTTATTACTATATTTCAGAGCAAACCTCC 3' |

**Suppl. Table ST5 - Yeast strains**

| Code | Genotype | Source |
|---|---|---|
| 501 | *h-, ura4-D18, ade6-704, leu1-32* | |
| 503 | *h+, ura4-D18, ade6-704, leu1-32* | |
| DHP260 | *h+, gcn5-Myc13::natMX6, ura-D18, ade6-M216* | Helmlinger et al., 2008 |
| DHP363 | *h+, gcn5-1(E191Q)-MYC13::natMX6, ura4-D18, ade6-M216* | Helmlinger et al., 2008 |
| DHP409 | *h-, Δsus1::URA4, ura-D18, ade6-M210* | Helmlinger et al., 2011 |
| NBY772 | *h-, nse4:GFP::HphMx4* | Pebernard et al., 2008 |
| MA1-6 | *h-, nse3-R254E::Lox, ura4-D18, ade6-704, leu1-32* | Zabrady et al., 2016 |
| MA23 | *h+, nse3-R254E::Lox, ura4-D18, ade6-704, leu1-32* | Zabrady et al., 2016 |
| MMP21 | *h-, nse4-FLAG-His::KanR leu1-32 ura4-D18* | Morikawa et al., 2004 |
| JMM956 | *h-, smc6-74, ade6-704, leu1-32, ura4-D18* | Verkade et al., 1999 |
| JMM1920 | *h-, smc6-X, ade6-704, leu1-32, ura4-D18* | Lehmann et al., 1995 |
| JMM2050 | *h+, nse2-SA::Ura4, leu1-32, ura4-d18,ade6-704* | Andrews et al., 2005 |
| PEM2 | *h-, mat1_m-cyhS, smt0, rpl42::cyhR (sP56Q), ade6-M210, leu1-32, ura4-D18* | Roguev et al., 2007 |
| Δada2 | *h+, Δada2::G418, ade6-216, ura4-D18, leu1-32* | BIONEER |
| Δada3 | *h+, Δada3::G418, ade6-216, ura4-D18, leu1-32* | BIONEER |
| Δbdc1 | *h+, Δbdc1::G418, ade6-210, ura4-D18, leu1-32* | BIONEER |
| Δbdf1 | *h+, Δbdf1::G418, ade6-216, ura4-D18, leu1-32* | BIONEER |
| Δcdt2 | *h+, Δcdt2::G418, ade6-216, ura4-D18, leu1-32* | BIONEER |
| Δdbl2 | *h+, Δdbl2::G418, ade6-216, ura4-D18, leu1-32* | BIONEER |
| Δdcn1 | *h+, Δdcn1::G418, ade6-216, ura4-D18, leu1-32* | BIONEER |
| Δddb1 | *h+, Δddb1::G418, ade6-216, ura4-D18, leu1-32* | BIONEER |
| Δeaf7 | *h+, Δeaf7::G418, ade6-216, ura4-D18, leu1-32* | BIONEER |
| Δgcn5 | *h+, Δ gcn5::G418, ade6-216, ura4-D18, leu1-32* | BIONEER |
| Δnth1 | *h+, Δnth1::G418, ade6-210, ura4-D18, leu1-32* | BIONEER |
| Δnup132 | *h+, Δnup132::G418, ade6-216, ura4-D18, leu1-32* | BIONEER |
| Δnup60 | *h+, Δnup60::G418, ade6-216, ura4-D18, leu1-32* | BIONEER |
| Δpng1 | *h+, Δpng1::G418, ade6-216, ura4-D18, leu1-32* | BIONEER |
| Δrad50 | *h+, Δrad50::G418, ade6-216, ura4-D18, leu1-32* | BIONEER |

| | | |
|---|---|---|
| Δrnh201 | *h+, Δrnh201::G418, ade6-216, ura4-D18, leu1-32* | BIONEER |
| Δsgf11 | *h+, Δsgf11::G418, ade6-216, ura4-D18, leu1-32* | BIONEER |
| Δsgf29 | *h+, Δsgf29::G418, ade6-210, ura4-D18, leu1-32* | BIONEER |
| Δsgf73 | *h+, Δsgf73::G418, ade6-216, ura4-D18, leu1-32* | BIONEER |
| ΔSPBC1347.08c | *h+, ΔSPBC1347.08c::G418, ade6-210, ura4-D18, leu1-32* | BIONEER |
| Δtra1 | *h+, Δtra1::G418, ade6-216, ura4-D18, leu1-32* | BIONEER |
| Δubp8 | *h+, Δubp8::G418, ade6-216, ura4-D18, leu1-32* | BIONEER |
| Δubr1 | *h+, Δubr1::G418, ade6-216, ura4-D18, leu1-32* | BIONEER |
| YLJ222 | *h-, NAT-nse3, mat1_m-cyhS, smt0, rpl42::cyhR (sP56Q), ade6-M210, leu1-32, ura4-D18* | This study |
| YLJ228 | *h-, NAT-nse3-R254E, mat1_m-cyhS, smt0, rpl42::cyhR (sP56Q), ade6-M210, leu1-32, ura4-D18* | This study |
| YLJ252 | *h-, Δsgf11::G418, nse3-R254E::Lox, ura4-D18, ade6-704?, ade6-M216?, leu1-32* | This study |
| YLJ255 | *h-, Δgcn5::G418, nse3-R254E::Lox, ura4-D18, ade6-704?, ade6-M216?, leu1-32* | This study |
| YLJ262 | *h-, Δsgf29::G418, nse3-R254E::Lox, ura4-D18, ade6-704?, ade6-M210?, leu1-32* | This study |
| YLJ265 | *h+, Δada2::G418, nse3-R254E::Lox, ura4-D18, ade6-704?, ade6-M216?, leu1-32* | This study |
| YLJ268 | *h-, Δubp8::G418, nse3-R254E::Lox, ura4-D18, ade6-704?, ade6-M216?, leu1-32* | This study |
| YLJ272 | *h-, Δngg1::G418, nse3-R254E::Lox, ura4-D18, ade6-704?, ade6-M216?, leu1-32* | This study |
| YLJ290 | *h+, Δsgf73::G418, nse3-R254E::Lox, ura4-D18, ade6-704?, ade6-M216?, leu1-32* | This study |
| YLJ375 | *h-, Δgcn5::G418, nse4-FLAG-His::KanR, ura4-D18, ade6-M216?, leu1-32* | This study |
| YLJ386 | *h-, Δgcn5::G418, ura4-D18, ade6-704?, ade6-M216?, leu1-32* | This study |
| YLJ388 | *h?, Δgcn5::G418, smc6-x, ura4-D18, ade6-704?, ade6-M216?, leu1-32* | This study |
| YLJ391 | *h?, Δgcn5::G418, smc6-74, ura4-D18, ade6-704?, ade6-M216?, leu1-32* | This study |
| YLJ426 | *h+, Δubp8::G418, nse4-FLAG-His::KanR, ade6-704?, ade6-M216?, leu1-32, ura4-D18?* | This study |
| YLJ459 | *h-, Δtra1::G418, nse3-R254E::Lox, ura4-D18, ade6-704?, ade6-M216?, leu1-32* | This study |
| YLJ462 | *h?, nse2-SA::URA4, Δgcn5::G418, ura4-D18, ade6-704?, ade6-M216?, leu1-32* | This study |
| YLJ496 | *h-, Δada2::G418, nse4-FLAG-His::KanR, ade6-704?, ade6-M216?, leu1-32, ura4-D18* | This study |
| YLJ507 | *h?, nse4-FLAG-His:KanR, gcn5-Myc13::natMX6, ura-D18, ade6-M216?, leu1-32?* | This study |
| YLJ510 | *h?, nse4-FLAG-His:KanR, gcn5-E191Q-Myc13::natMX6, ura-D18, ade6-M216?, leu1-32?* | This study |
| YPK150 | *h?, nse4:GFP::HphMx4, Δgcn5::G418, ade6-216?, ura4-D18?, leu1-32?* | This study |
| YPK165 | *h?, nse4:GFP::HphMx4, Δbrc1::G418, leu1-32?, ura4-D18?* | This study |

**Suppl. Table ST6 - qPCR primers**

| | | | | |
|---|---|---|---|---|
| *vht1* | I:4959888-4960015 | prBS103 | fw | 5′ GAGGAGACAAAGTTTTCATCCATTTGGG 3′ |
| | | prBS104 | rev | 5′ GGGAGTCGTTGCTATTGCCC 3′ |
| *fip1* | I:4233546-4233678 | prBS105 | fw | 5′ CGACGTTCCTTGTACCACATGAGG 3′ |
| | | prBS106 | rev | 5′ GGTGGTTACATGATCTTCAATGCCA 3′ |
| *rps502* | I:3496612-3496743 | prBS113 | fw | 5′ GGAAAGCAGCCTCACGAGC 3′ |
| | | prBS114 | rev | 5′ GTCCTCGTGAAGACTCCACTCG 3′ |
| *ser3* | III:470819-470947 | prBS109 | fw | 5′ CGATAAAAAGCTCGTTAACTTGGCG 3′ |
| | | prBS110 | rev | 5′ GGTGCTGTTAACTTCCCTGAGG 3′ |
| *tef3* | III:1688122-1688266 | prBS111 | fw | 5′ GAGTGCTCTTTCCTCGTCGG 3′ |
| | | prBS112 | rev | 5′ CTCAGCACGGTCAACTTCAGC 3′ |
| *slx9* | I:499186-499310 | prBS129 | fw | 5′ CGGTAAATTGATACGCCC 3′ |
| | | prBS130 | rev | 5′ GACATCCCGACAATACATC 3′ |
